# A dynamical limit to evolutionary adaptation

**DOI:** 10.1101/2023.07.31.551320

**Authors:** Matthew J. Melissa, Michael M. Desai

## Abstract

Natural selection makes evolutionary adaptation possible even if the overwhelming majority of new mutations are deleterious. However, in rapidly evolving populations where numerous linked mutations occur and segregate simultaneously, clonal interference and genetic hitchhiking can limit the efficiency of selection, allowing deleterious mutations to accumulate over time. This can in principle overwhelm the fitness increases provided by beneficial mutations, leading to an overall fitness decline. Here, we analyze the conditions under which evolution will tend to drive populations to higher versus lower fitness. Our analysis focuses on quantifying the boundary between these two regimes, as a function of parameters such as population size, mutation rates, and selection pressures. This boundary represents a state in which adaptation is precisely balanced by Muller’s ratchet, and we show that it can be characterized by rapid molecular evolution without any net fitness change. Finally, we consider the implications of global fitness-mediated epistasis, and find that under some circumstances this can drive populations towards the boundary state, which can thus represent a long-term evolutionary attractor.

## I. INTRODUCTION

Evolution is often thought of as an optimization process, in which natural selection pushes populations inevitably uphill, towards a local optimum in the fitness landscape (Wright, 1932). However, much recent work has shown that in many populations, numerous linked mutations often arise and segregate simultaneously (Barroso-Batista et al., 2014; De Visser et al., 1999; Desai et al., 2007; Kao and Sherlock, 2008; Lang et al., 2013; Miralles et al., 1999; Nourmohammad et al., 2019; Strelkowa and Lässig, 2012). In these *rapidly evolving* populations, natural selection is much less efficient: it cannot act on each mutation independently (Gerrish and Lenski, 1998). As a result, deleterious mutations can often fix, which can slow down adaptation or even reverse its direction, leading to declining fitness over time.

Extensive previous work has studied the accumulation of deleterious mutations via Muller’s ratchet (Felsenstein, 1974; Muller, 1964), particularly in models in which beneficial mutations are either negligible or can be treated as a rare perturbation (Bachtrog and Gordo, 2004; Neher and Shraiman, 2012; Rouzine et al., 2008). Similarly, numerous studies have considered the accumulation of beneficial mutations (i.e. adaptation) when deleterious mutations are absent (Desai and Fisher, 2007; Fisher, 2013; Good et al., 2012) or can be treated as a perturbation (Good and Desai, 2014; Johnson and Barton, 2002). However, we lack an understanding of the interplay between the accumulation of beneficial and deleterious mutations more generally. Except in special cases (e.g. when clonal interference is absent (Whitlock, 2000), or when all beneficial and deleterious mutations have the same fitness effect (Goyal et al., 2012)), this has made it impossible to answer a very basic question: given a particular set of population genetic parameters (population size, mutation rate, and fitness landscape), will a population tend to increase or decrease in fitness? In other words, under what circumstances can evolution act as an optimization process, and when do populations actually move towards less-optimal genotypes?

Here, we analyze this interplay between beneficial and deleterious mutations in rapidly evolving populations, in the regime where both types of mutations can be important. Our analysis leverages recent work in traveling wave models of evolutionary dynamics, and in particular our recently introduced *moderate selection, strong-mutation* (MSSM) approximation (Melissa et al., 2022). Using this approach, we predict the conditions under which populations will tend to increase or decrease in fitness (i.e. where the rate of change in mean fitness, *v*, is positive or negative). The boundary surface between these two regions of the parameter space, at which *v* = 0, corresponds to a state in which beneficial and deleterious mutations accumulate in a balanced way. While the fitness trajectory of a population in the *v* = 0 state appears neutral, the evolutionary dynamics of these populations can be strongly nonneutral. For example, a steady state accumulation of weakly deleterious mutations may be offset by the fixation of beneficial mutations under moderate or strong selection. We also consider additional surfaces of the parameter space on which patterns of molecular divergence and genetic diversity would suggest a population has evolved neutrally or nearly neutrally, but in fact mask a balance between the competing signatures of positive and negative selection.

We conclude by considering how our results and the structure of the fitness landscape determine the long-term outcomes of evolution. For example, it is natural to expect that beneficial mutations become less common (and deleterious mutations more common) as a population increases in fitness. This will tend to lead a population not towards a local optimum, but instead towards the *v* = 0 state (see e.g. Goyal et al. (2012)). More generally, recent empirical work has identified a consistent pattern of diminishing returns epistasis: beneficial mutations tend to have weaker effects as populations increase in fitness (Bakerlee et al., 2021; Chou et al., 2011; Kryazhimskiy et al., 2014). An analogous pattern for epistasis on deleterious mutations is less clear, but recent work has identified a trend in which deleterious mutations are more costly in more-fit backgrounds (Johnson and Desai, 2022; Johnson et al., 2019). We show here that, depending on details of the landscape and the starting point, these and other patterns of fitness-mediated epistasis can often (but not always) drive a population towards the *v* = 0 state. Thus the *v* = 0 state can in some circumstances represent a long-term evolutionary attractor, and define the extent to which evolution can act to optimize fitness.

## II. MODEL

We model the evolution of a population of N haploid individuals, in which random mutations arise within a specific genomic region at a total rate U. We assume that recombination can be neglected within this region on the relevant timescales. We assume that each new mutation confers a fitness effect, s, drawn from some distribution of fitness effects (DFE), 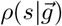, that depends on the genotype 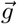 of an individual as well as the environment the population evolves in (its fitness landscape). The DFE includes both beneficial and deleterious mutations, with beneficial mutations corresponding to s > 0 and deleterious mutations corresponding to s < 0. To be more precise, a mutation with effect s increments an individual’s (log) fitness X by an amount s, and we assume offspring numbers are drawn from a multinomial distribution each generation; the expected offspring number of an individual with fitness *X* is 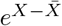, where 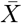 denotes the mean fitness of the population.

The genotype-dependence of 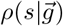 has been termed *macroscopic epistasis* (Good and Desai, 2015). This macroscopic epistasis can arise due to individual *microscopic epistatic* interactions among specific mutations, which collectively determine the overall DFE for a given genotype. We make the key assumption that similar genotypes share a similar 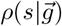, and in particular, that those genotypes simultaneously present in a population (which are similar because of their relatedness by common ancestry) share the same DFE, *ρ*(*s*). This allows us to solve for the dynamics by treating *ρ*(*s*) instantaneously as a constant parameter. We note that this assumption can be satisfied even in the presence of pervasive microscopic epistasis, as long as idiosyncratic interactions among mutations largely “average out” in contributing to the full distribution 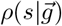.

For simplicity, we focus on a few simplifying forms of *ρ*(*s*) in our analysis. For instance, we consider the case where all beneficial mutations have effect *s*_*b*_ and all deleterious mutations have effect −*s*_*d*_ (with *s*_*b*_, *s*_*d*_ > 0 by convention). This example is useful for building general intuition, and is motivated by recent work showing that the evolutionary dynamics of rapidly evolving populations can in many cases be well-captured by a DFE consisting of a single appropriately-chosen “predominant” effect size (Fisher, 2013; Good et al., 2012; Hegreness et al., 2006). We also consider the cases of exponentially-distributed (and more generally gamma-distributed) effects of beneficial and of deleterious mutations, though our analysis can be extended to more general DFEs relatively straightforwardly. Importantly, we make no assumption that the DFEs of beneficial mutations and of deleterious mutations are the same or similar in shape or in scale.

## III. RESULTS

The central goal of our analysis is to determine whether a population will tend to increase or decrease in fitness for a given set of parameters: the population size, *N*, the mutation rate, *U*, and the distribution of fitness effects, *ρ*(*s*). Because our goal is to determine whether *v* > 0 or *v* < 0 for a given set of parameters, we focus on analyzing the boundary between these two regimes. This boundary is by definition a *v* = 0 surface where the mean fitness of the population does not on average either increase or decrease. We will find it useful to write the average rate of change in mean fitness, *v*, in terms of the fixation probability of a new mutation, *p*_fix_(*s*),

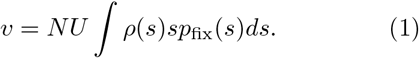

To find the *v* = 0 surface we then set Eq. (1) equal to 0, which gives a constraint on the parameters *N, U*, and *ρ*(*s*) that defines the *v* = 0 surface in parameter space.

We can clearly have *v* = 0 if selection on deleterious mutations is sufficiently strong and beneficial mutations are sufficiently rare that no selected mutations fix at all, and the evolutionary dynamics are entirely neutral (i.e. if *ρ*(*s*)p_fix_(*s*) is negligible for all *s*). Apart from this trivial case, the *v* = 0 state by definition involves substantial accumulation of deleterious mutations (at least relative to the accumulation of beneficial mutations) which can be facilitated by the effects of linked selection and clonal interference. For instance, deleterious mutations may routinely hitchhike along with, or hinder the fixation of, a beneficial mutation (Charlesworth, 1994; Peck, 1994; Smith and Haigh, 1974). Interference among multiple beneficial mutations may also substantially reduce the rate at which they can fix in the population (Gerrish and Lenski, 1998; Hill and Robertson, 1966). To obtain an accurate description of the *v* = 0 state, our expression for the fixation probabilities *p*_fix_(*s*) must therefore take these effects into account.

Frequent interference among mutations is a defining feature of *rapid evolution*, which has been the focus of much recent theoretical work (Good and Desai, 2014; Good et al., 2012; Hallatschek, 2011; Neher and Hallatschek, 2013; Neher et al., 2010; Rouzine et al., 2008, 2003). Broadly speaking, this work uses traveling wave models, which first analyze the steady-state distribution of fitness within the population (the “traveling wave of fitness”), and then use this as the basis for computing the fixation probabilities of new mutations, *p*_fix_(*s*), and the average rate of fitness increase or decline, v. Most work on traveling wave models has been done by considering only beneficial mutations (Desai and Fisher, 2007; Fisher, 2013; Good et al., 2012) or only deleterious mutations (Neher and Shraiman, 2012), or by focusing on one type and treating the other perturbatively (Good and Desai, 2014). A key exception is the *moderate selection, strong-mutation* (MSSM) approximation we have recently introduced (Melissa et al., 2022), which can be applied to analyze rapidly evolving populations for which both beneficial and deleterious mutations affect the dynamics in a substantial way. Here, we use this MSSM approximation to analytically describe the *v* = 0 state.

A key result is that within the MSSM regime (which we discuss below), the fixation probability of a new mutation is given by

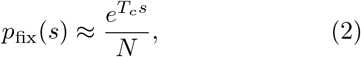

valid for both positive and negative s. Here *T*_*c*_ is a derived quantity whose definition and relationship to the parameters *N, U* and *ρ*(*s*) we reproduce in *SI Appendix*, and which approximately equals ⟨*T*_2_⟩ /2— one-half the average time since two randomly chosen individuals share a common ancestor (i.e., a *coalescence* timescale). We note that in much of the population genetics literature, the average pair-wise coalescent time ⟨*T*_2_⟩ is identified with an effective population size *N*_*e*_. We avoid this language here because in general, the evolutionary dynamics in rapidly evolving populations are not equivalent to those in a neutrally evolving population for any choice of *N*_*e*_ (Neher, 2013).

Eq. (2) differs from the standard formula for the fixation probabilities of independently evolving loci (Crow et al., 1970), which in our notation can be written as

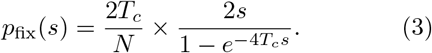

Eq. (3) has been used by Whitlock (2000) to address similar questions, although that work treats *T*_*c*_ (referred to as *N*_*e*_/2) as an independent parameter, instead of considering how it depends on the population parameters *N, U*, and *ρ*(*s)*. For the sake of comparison, we discuss the predictions following from Eq. (3) alongside our results below. The predictions are qualitatively (and even quantitatively) similar in some respects, but they break down in other cases. This is unsurprising in light of recent work that has shown that Eq. (3) fails to adequately describe the fixation probabilities of mutations in the presence of widespread linked selection, particularly when mutations confer fitness effects on a wide range of scales (Messer and Petrov, 2013).

Eq. (2) immediately implies that if we scale fitness effects to the coalescence timescale by defining γ≡ T_*c*_*s*, the *v* = 0 surface is defined by the concise equation

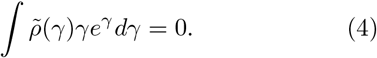

Eq. (4) implies that we can characterize the *v* = 0 surface given only the distribution of “scaled” fitness effects, 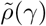 (as well as validity of the MSSM approximation, which we discuss below). We emphasize that this is not by itself sufficient to determine how the *v* = 0 surface depends on the underlying parameters, because *T*_*c*_ depends in a nontrivial way on *N, U*, and *ρ*(*s*). We return to this dependence in more detail below. However, in the next section we first analyze key properties of the *v* = 0 surface in the space of scaled selective effects, focusing particular attention on two specific choices of the DFE as representative examples.

### A. The *v* = 0 surface in the space of scaled effects

In the scaled parameter space, the *v* = 0 surface depends only on 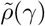, and not on the population size *N* or the mutation rate *U* (which enter only through their effect in determining *T*_*c*_). To gain qualitative insight, we begin by considering the simple case in which all beneficial mutations confer a single scaled effect, γ_*b*_≡ *T*_*c*_*s*_*b*_, and all deleterious mutations confer a (potentially different) single scaled effect, −γ_*d*_ ≡ −*T*_*c*_s_*d*_ (with s_*b*_, s_*d*_ > 0 by convention). Specifically, we have 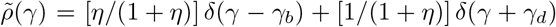, where we have defined η≡ *U*_*b*_/*U*_*d*_ as the ratio of beneficial to deleterious mutation rates. Plugging this into Eq. (4), we find that within the 3-dimensional parameter space spanned by *γ*_*b*_, *γ*_*d*_, and η, the *v* = 0 constraint is a 2-dimensional surface given by

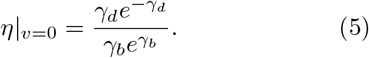

In Fig. 1 we validate this prediction for the *v* = 0 surface. To do so, we conducted Wright-Fisher simulations for populations whose parameters lie on a grid with varying η, *Ns*_*b*_ and *Ns*_*d*_. Each simulated population is plotted using its corresponding value of *T*_*c*_, measured by observing its pairwise neutral heterozygosity π_neu_ averaged over simulation runs (with *T*_*c*_ taken as ⟨*T*_2_⟩ /2 = π_neu_/(4*U*_*n*_), where *U*_*n*_ is the neutral mutation rate used in simulations). The prediction in Eq. (5) qualitatively (and except perhaps for γ_*d*_ ≫ 1, quantitatively) describes the *v* = 0 surface in the space of scaled fitness effects γ_*b*_ and γ_*d*_. The simulations represented in Fig. 1 are all conducted for populations with *NU* = 10^4^. In Fig. S1, we present the results of additional simulations which include the cases *NU* = 10^3^ and *NU* = 10^2^; similar agreement is obtained.

**FIG. 1.**
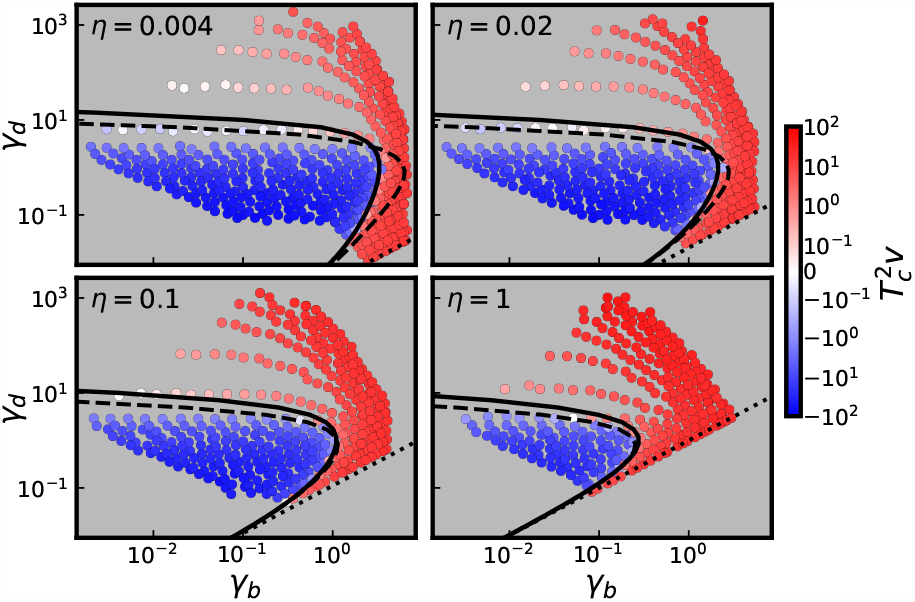
Cross sections of the *v* = 0 surface in the space of scaled effects *γ*_*d*_ vs. *γ*_*b*_, for the four values of *η* denoted above. Each point corresponds to a simulated parameter combination, colored by its measured value of 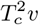. Solid line denotes the prediction for the *v* = 0 curve given by Eq. (4). Dashed lines denotes the prediction for the *v* = 0 curve obtained using Eq. (3). Dotted lines denote the lines on which *η* = *γ*_*d*_*/γ*_*b*_. Simulated parameter combinations lie on the grids of logarithmically spaced *Ns*_*d*_ and *Ns*_*b*_ values depicted in Fig. 2, with DFEs consisting of a single beneficial effect and a single deleterious effect, and with *NU* = 10^4^.

Several qualitative features of Eq. (5) are notable. If the selective effects of both beneficial and deleterious mutations are small compared to 1/*T*_*c*_ (i.e. γ_*b*_ ≪ 1 and γ_*d*_ ≪ 1), we see from Eq. (5) that the *v* = 0 surface is defined by *η*|_*v*=0_≈ *γ*_*d*_/*γ*_*b*_ (dotted line in Fig. 1; note this converges with the *v* = 0 surface observed in simulations when γ_*b*_ ≪ 1 and γ_*d*_ ≪ 1). This corresponds to *U*_*b*_*s*_*b*_ = *U*_*d*_*s*_*d*_, the surface on which *v* = 0 if beneficial and deleterious mutations accumulate neutrally, such that each mutation fixes with probability 1/*N*. The surface *η* = *γ*_*d*_/*γ*_*b*_ can be thought of as an upper bound to the actual *v* = 0 surface; as the strength of selection (i.e. *γ*_*b*_ and/or *γ*_*d*_) is increased, the actual fraction η required to have *v* = 0 will always be smaller than γ_*d*_/γ_*b*_.

More generally, if we increase γ_*b*_ at fixed γ_*d*_, *η*|_*v*=0_ decreases: a smaller ratio η of beneficial to deleterious mutations is required to be in the *v* = 0 state. This makes intuitive sense: increasing *γ*_*b*_ increases both the fixation probability of beneficial mutations and the fitness benefit they provide to the population upon fixing. The effect of changing *γ*_*d*_ is more subtle, because while increasing *γ*_*d*_ decreases the fix-ation probability of deleterious mutations, it also increases the fitness cost to the population they incur upon fixing. This means that the expected fitness costs to the population of deleterious mutations are not monotonic with effect size: for a fixed value of γ_*b*_ (below a threshold value described below), the population will adapt for small *γ*_*d*_, decline in fitness for intermediate *γ*_*d*_, and adapt for large *γ*_*d*_. This is because for sufficiently small *γ*_*d*_, deleterious mutations fix routinely but do not confer large enough effects to counteract the beneficial mutations which fix, while for sufficiently large *γ*_*d*_, deleterious mutations are purged by selection too efficiently to counteract the fixation of beneficial mutations. Instead, deleterious mutations are maximally impactful (in the sense that *η*|_*v*=0_ is maximized) at the intermediate scaled effect size 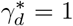. For sufficiently large *γ*_*b*_ and/or η, deleterious mutations cannot lead to decline in fitness for *any* value of *γ*_*d*_ (although *T*_*c*_v will still be minimized at *γ*_*d*_ = 1). For example, at a given η, the population will always adapt provided that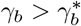, with

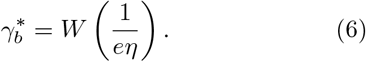

Here *W* (*x*) denotes the Lambert W function satisfying *W*(*x*)*e*^*W* (*x*)^ = *x*, which has the asymptotic behavior *W* (*x*) ∼ log *x* − log log *x* for large *x*. We can think of the curve 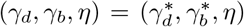, parameterized by *η*, as a “ridgeline” of the *v* = 0 surface, on which *η* is maximized as a function of *γ*_*d*_.

The above analysis can be extended straightforwardly to a full distribution of fitness effects. As a simple example, we consider the case in which both beneficial and deleterious mutations are drawn from exponential distributions with mean scaled effects *γ*_*b*_ and *γ*_*d*_ respectively, and a ratio *η* = *U*_*b*_/*U*_*d*_ of beneficial to deleterious mutations (in the *SI Appendix*, we extend these results to the case of gamma-distributed DFEs, and comment further on arbitrary DFEs). Specifically, we have 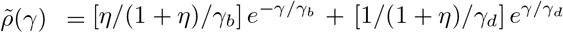. Plugging this scaled DFE into Eq. (4), we find that the *v* = 0 surface is defined by

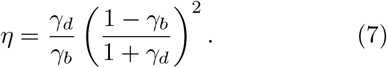

We note that in this case, a requirement that *γ*_*b*_ < 1 dynamically emerges given validity of the MSSM approximation (and thus the *η* given by Eq. (7) is positive for all relevant *γ*_*b*_ and *γ*_*d*_) while *γ*_*d*_ > 1 is permitted. As in the case of a two-effect DFE, a *v* = 0 “ridgeline” exists on which *η* is maximized as a function of *γ*_*d*_, and on which 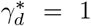 and 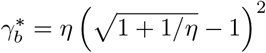. The behavior of the *v* = 0 surface for large *γ*_*d*_ is qualitatively different, however. Instead of the dependence *η* ∝ *γ*_*d*_*e*^−*γd*^, we have the much slower falloff *η* ∝ 1/γ_*d*_ for large *γ*_*d*_. The dependence 1/*γ*_*d*_ in turn approximates the fraction of deleterious effects with |*γ*| < 1—that is, the fraction of deleterious effects with a reasonable chance of fixing—as opposed to the fixation probability ∝*e*^−*γd*^ of the average deleterious effect. As a result, compared to the case of a two-effect DFE, *v* = 0 curves for a full DFE have a much more broad decay of *γ*_*b*_ toward zero at large *γ*_*d*_. In Fig. S1 we compare the prediction in Eq. (7) to the results of simulations for the same *η* values considered in Fig. 1, and for *NU* values ranging from 10^2^ to 10^4^. Our results are qualitatively similar to the case of single fitness effects shown in Fig. 1, although agreement begins to break down—particularly for 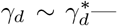 for smaller *NU* and smaller *η* (i.e. for smaller values of *NU*_*b*_).

### B. Computing *T*_*c*_ in terms of the population genetic parameters, *N* and *Uρ*(*s*)

Above, we described the *v* = 0 constraint on the distribution of *scaled* effects *γ* = *T*_*c*_s. Expressed in this way, the *v* = 0 constraint takes on the simple analytical form in Eq. (4). In certain cases, the distribution of scaled effects *γ* is more readily probed than the distribution of *unscaled* effects s. For instance, given DNA sequencing data from a population at a fixed point in time (e.g. an observation of its site frequency spectra, of both synonymous mutations and nonsynonymous mutations), its distribution of scaled fitness effects can be inferred (Eyre-Walker and Keightley, 2007; Keightley and Eyre-Walker, 2010; Sawyer and Hartl, 1992). To infer the distribution *unscaled* fitness effects s then requires an independent estimate of the coalescence timescale, which is typically confounded with estimates of the neutral mutation rate (Charlesworth, 2009). However, in certain cases—such as in the context of experimentally evolved populations (Frenkel et al., 2014; Levy et al., 2015), the distribution of unscaled effects *s* may be more practical to measure. For this reason, it may be more useful to obtain a *v* = 0 constraint on the parameters *N* and *Uρ*(s); this can also be useful in building an intuitive understanding for how shifts in these underlying parameters affect adaptation or fitness decline. As we will see in the Discussion, a *v* = 0 constraint on unscaled effects s may also be more useful in working out the long-term implications of simple patterns of fitness-mediated epistasis observed empirically, for which the parameter *Uρ*(s) may vary in a more straightforward way, over the course of an evolutionary trajectory, than the distribution of scaled effects.

In general, the coalescence timescale *T*_*c*_ depends in a complicated way on the parameters *N* and *Uρ*(s), and except in special cases, the mapping between a distribution of unscaled effects *Ns* and the distribution of scaled effects *T*_*c*_*s* is not particularly clear. The MSSM approximation yields a relation between *T*_*c*_ and the underlying parameters *N* and *Uρ*(*s*)— and can thus be used to obtain a *v* = 0 constraint on the parameters *N* and *Uρ*(*s*). Importantly, this can be contrasted to a prediction for the *v* = 0 surface using Eq. (3), which is not associated with any relation between *T*_*c*_ and the underlying parameters. We reproduce the relation between *T*_*c*_ and *N* under the MSSM approximation in the *SI Appendix*. The key result is

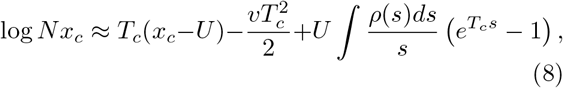

where we have defined

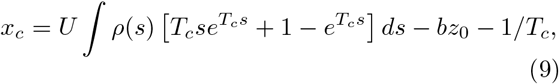

with

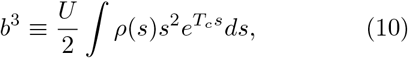

and where *z*_0_≈− 2.34 is the least negative zero of the Airy function Ai(*z*). This relation is an implicit equation for *T*_*c*_, which precludes writing down a simple analytical form of the *v* = 0 constraint such as that in Eq. (5). However, it is straightforward to solve Eq. (8) for *T*_*c*_ in terms of *N* numerically (more precisely, we solve a less approximated version of Eq. (8); see *SI Appendix* for details). We can then plug this result into Eq. (4) to obtain a condition for the *v* = 0 surface in terms of the underlying population genetic parameters, *N, U*, and *ρ*(*s*).

### C. Validity of the MSSM approximation

The results for *p*_fix_(*s*) in Eq. (2) and particularly for *T*_*c*_ in Eq. (8) both depend on the validity of the MSSM approximation. We review the conditions of validity of the MSSM approximation in the *SI Appendix*. These conditions can be concisely expressed in terms of *T*_*c*_, the range Δ*x*_*f*_ of fitness advantages from which an eventual common ancestor of the population is typically supplied (which is well-approximated by b as defined above), and the sizes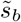 and 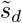of typical fixed beneficial and deleterious effects. Roughly speaking, the MSSM approximation requires that *T*_*c*_Δ*x*_*f*_ ≫1 (which ensures the population is rapidly evolving and that the dynamics by which mutations fix are strongly nonneutral) and that 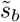 and 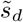 are both small compared to Δ*x*_*f*_ ; in addition, the distribution of beneficial fitness effects must fall off exponentially, or faster than exponentially, with large s. Given the solution for *T*_*c*_ from Eq. (8), it is straightforward to determine whether these conditions are met for a particular point on the *v* = 0 surface.

The points 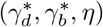 lying along the *v* = 0 “ridge-line” are convenient landmarks of *v* = 0 curves at which the validity of the MSSM approximation can be assessed. At these points, selection is sufficiently strong that the MSSM predictions are nontrivial, and the *v* = 0 surface is guaranteed to deviate substantially from the surface *η* = *γ*_*d*_/γ_*b*_ on which *v* = 0 assuming neutral accumulation. Furthermore, validity of the MSSM approximation at 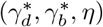 implies validity of the approach for smaller values of *γ*_*d*_ and *γ*_*b*_ along the same *v* = 0 curve, up to and including 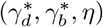, for a given *η*. Note that in the limit *γ*_*b*_ → 0 and *γ*_*d*_ → 0, the population dynamics become purely neutral, and the condition *T*_*c*_Δ*x*_*f*_ ≫ 1 required of the MSSM approximation breaks down; however, by this point, the *v* = 0 surface will simply approach its neutral expectation *η* = *γ*_*d*_/*γ*_*b*_.

In the *SI Appendix*, we assess the validity of the MSSM approximation at the points 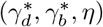, for the cases of a two-effect DFE and a two-exponential DFE. We find that in both of these cases, the conditions of validity of the MSSM approximation are met provided that *T*_*c*_*U*_*b*_ ≫ 1 at 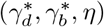; note that we assume *U*_*b*_ ≤ *U*_*d*_ throughout, so that *T*_*c*_*U*_*d*_ ≫ 1 is also implied. This condition still makes reference to the quantity *T*_*c*_, which can be determined using Eq. (8). However, the condition *T*_*c*_*U*_*b*_≫ 1 has a relatively simple dynamical interpretation—essentially, that a given lineage will acquire multiple beneficial mutations over the coalescence timescale—and is satisfied at 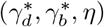for sufficiently large *NU* (since *T*_*c*_*U* increases with *NU*, if *γ*_*b*_, *γ*_*d*_ and *η* are held fixed). This suggests that the MSSM approximation is of broad use in the describing the 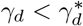 portion of *v* = 0 curves for rapidly evolving populations in which interference is important.

While the MSSM approximation is expected to be valid in describing the 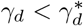 portion of the *v* = 0 surface, it may break down in the presence of strong deleterious effects 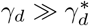, even if *T*_*c*_*U*_*b*_ ≫ 1. Here, however, we will see that an alternative heuristic approach works well: we take

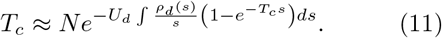

The form of *T*_*c*_ in Eq. (11) can be motivated by analyzing how strongly deleterious mutations contribute to the determination of *T*_*c*_ in the MSSM approximation (that is, through the final term in Eq. (8)). We note that in using Eq. (11) we neglect any effect of beneficial mutations on the determination of *T*_*c*_, which is reasonable when beneficial effects are sufficiently weak and deleterious effects sufficiently strong. When deleterious mutations all confer a single deleterious effect with *T*_*c*_*s*_*d*_ ≫ 1, *T*_*c*_ in Eq. (11) reduces to the well-known quantity 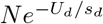. This is often referred to as *N*_*e*_, and under certain conditions gives the number of deleterious-mutation-free individuals in the population at equilibrium (Haigh, 1978), and in turn the coalescence timescale (Charlesworth et al., 1993). A similar interpretation can be given if *T*_*c*_*s* ≫ 1 for all possible deleterious effects (Good et al., 2014; Söderberg and Berg, 2007). When instead both weak and strong deleterious effects are possible, the factor 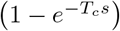 essentially picks out those deleterious effects with *T*_*c*_*s* ≫ 1 (which are important in reducing the effective population size and related coalescence timescale). Here, we refer to this heuristic approach to estimating *T*_*c*_ as an “*N*_*e*_-based heuristic”; by substituting this *T*_*c*_ into Eq. (3) we can obtain predictions for *v* = 0 curves in the same way *v* = 0 curves are obtained above using the MSSM approximation.

### D. The *v* = 0 constraint in terms of population genetic parameters

We compare our predictions for the *v* = 0 constraint to the results of simulations in Fig. 2. We can see that the small *Ns*_*d*_ and large *Ns*_*d*_ portions of *v* = 0 curves are well-described by the MSSM approximation and our *N*_*e*_-based heuristic, respectively. For concreteness, we connect these two approaches by taking the result of the MSSM approximation for *Ns*_*d*_ up to and including (*Ns*_*d*_)^*^, the maximal value of *Ns*_*d*_ on a *v* = 0 curve, given *η* and *NU*. We therefore consider the following prediction for *v* = 0 curves:

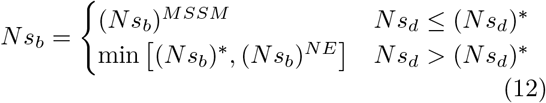

where (*Ns*_*b*_)^*MSSM*^ and (*Ns*_*b*_)^*NE*^ denote predictions for *Ns*_*b*_ along *v*= 0 curves, given the MSSM approximation and *N*_*e*_-based heuristic, respectively; (*Ns*_*d*_)^*^ and (*Ns*_*b*_)^*^ are predictions for maximal *Ns*_*d*_ value and corresponding *Ns*_*b*_ at which v = 0, obtained using the MSSM approximation. In Fig. 2 we can see that Eq. (12) adequately predict *v* = 0 curves across a range of *Ns*_*d*_ values and *η* values for both the cases of two-effect and two-exponential DFEs. These simulations are conducted with *NU* = 10^4^; the results of simulations with *NU* = 10^2^ and *NU* = 10^3^ are presented in Fig. S2. As is observed for *v* = 0 curves in the space of scaled effects, we note that the *v* = 0 curves are much less “sharp” at large values of *Ns*_*d*_ for the cases of two-exponential DFEs, as compared to cases of two-effect DFEs. Note that because at fixed *η* and *NU, T*_*c*_ varies with *Ns*_*b*_ and *Ns*_*d*_, the point in parameter space at which 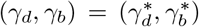 is not the same as the point at which (*Ns*_*b*_, *Ns*_*d*_) = ((*Ns*_*b*_),(*Ns*_*d*_)^*^). However, we can see in Fig. 2 and Fig. S2—with the point at which 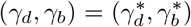 denoted by a star marker—that for all cases considered, these points nearly coincide. Thus, validity of the MSSM approximation up to one of these points essentially implies validity up to the other point, which motivates us to use the predictions of the MSSM approximation for *Ns*_*d*_ up to (*Ns*_*d*_)^*^ in Eq. (12).

**FIG. 2.**
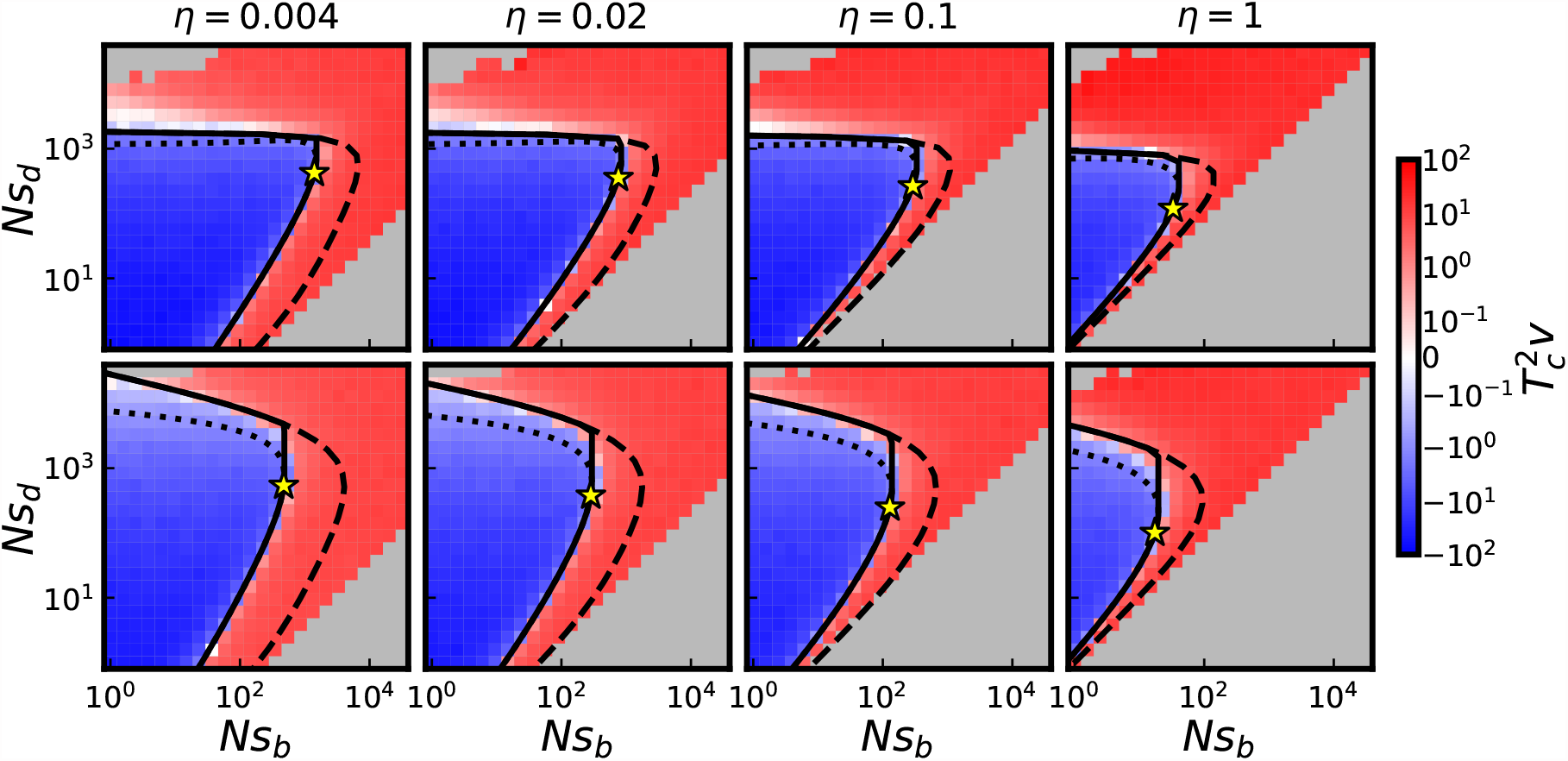
Cross-sections of the *v* = 0 surface in the space of unscaled effects *Ns*_*d*_ vs. *Ns*_*b*_, for *NU* = 10^4^ and four values of *η*, with populations colored by simulated values of 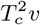. Populations in the top row are subject to a two-effect DFE; populations in the bottom row are subject to a two-exponential DFE. Solid lines denote predictions of the *v* = 0 surface obtained by connecting predictions obtained using the MSSM approximation and *N*_*e*_-based heuristic (that is, using Eq. (12)). Dotted lines denote predictions obtained using the MSSM approximation, for *Ns*_*d*_ *>* (*Ns*_*d*_)^*^; dashed lines denote predictions of the *N*_*e*_-based heuristic. Stars denote points at which 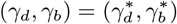, as computed by MSSM approximation. Gray squares denote parameter combinations not simulated because *U*_*d*_*s*_*d*_ *< U*_*b*_*s*_*b*_, or parameter combinations eliminated from consideration because fewer than 10 epochs were reached during the simulation runtime.

To summarize the accuracy of the MSSM approximation in predicting *v* = 0 curves across the entire range of *NU* and *η* values simulated, in Fig. 3 we compare our predictions for (*Ns*_*b*_)^*^, (*Ns*_*d*_)^*^, 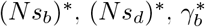and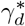 to the corresponding quantities obtained from simulations (with details of how these quantities are extracted from simulations provided in *Materials and Methods*). Fig. 3 includes populations which are subject to two-effect DFEs; we provide the same comparison for populations instead subject to two-exponential DFEs in Fig. S3. We can see that these quantities (and thus the weaker-selection portion of *v* = 0 curves) are well-predicted as long as *T*_*c*_*U*_*b*_ ≫ 1 at a given point along the *v* = 0 ridgeline.

**FIG. 3.**
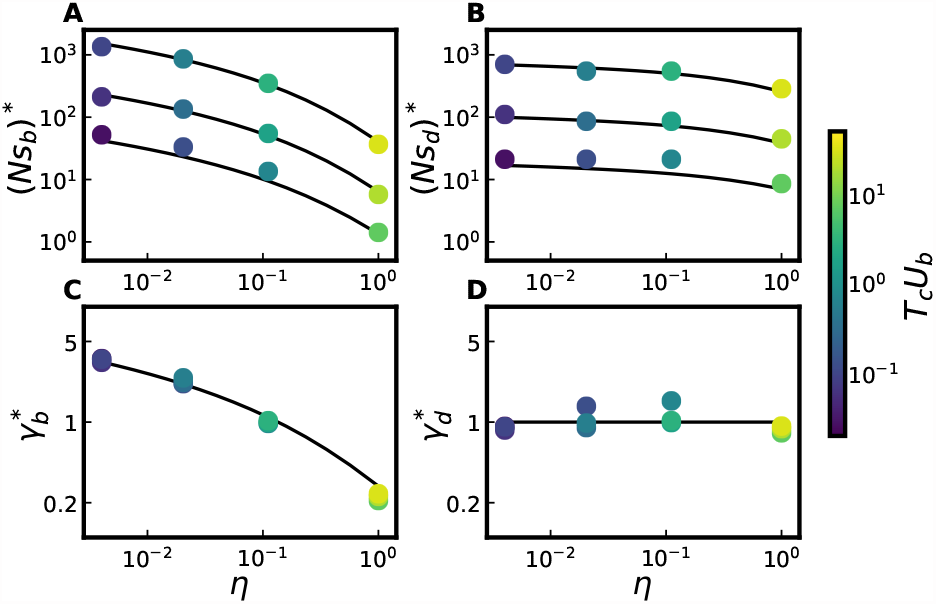
Comparison between simulations and MSSM predictions for, in panels A and B, the extremal point ((*Ns*_*b*_)^*^, (*Ns*_*d*_)^*^) and, in panels C and D, the ridgeline point 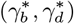. Each point is obtained from the simulation results depicted in a particular panel of Fig. 2 or Fig. S2 (with values of *η* denoted on the horizontal axis and values of *NU*∈ { 10^2^, 10^3^, 10^4^}, and with populations subject to a two-effect DFE); for details on how ridgeline and extremal points are extracted from a given panel of simulation results, see *Materials and Methods*. In panels A and B, the top theory curve corresponds to *NU* = 10^4^, the middle theory curve corresponds to *NU* = 10^3^, and the bottom theory curve corresponds to *NU* = 10^2^. Points are colored according to their values of *T*_*c*_*U*_*b*_ at 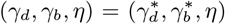, with *T*_*c*_ measured in simulations through levels of pairwise neutral heterozygosity.

In Fig. 2, we have chosen to illustrate cross sections of the parameter space spanned by the axes *Ns*_*b*_ and *Ns*_*d*_ because of the striking nonmonotonicity of *v* = 0 curves in this space (at fixed *η* and *NU*). To gain a more complete qualitative understanding of the *v* = 0 constraint, we can consider small perturbations of parameters from the *v* = 0 constraint along each of the two remaining axes, *η* and *NU*. The behavior with *η* is simple: *N* ^2^*v* increases monotonically with *η* (with other parameters held fixed), so *v* = 0 curves are shifted (and distorted, to some extent) towards *lower Ns*_*b*_ values as *η* is increased (i.e. with larger η, smaller values of *Ns*_*b*_ are needed to have *v* = 0). The behavior with *NU* is less immediately clear, but also straightforward: *v* = 0 curves are shifted toward *larger Ns*_*b*_ values as *NU* is increased. This reflects the fact that a larger *NU* value implies more frequent interference, and thus less efficient selection for beneficial mutations and against deleterious mutations. From this dependence, the behavior with *U* (at fixed η, *N*, s_*b*_ and s_*d*_) also follows: *v* = 0 curves are shifted toward larger *Ns*_*b*_ values as *U* is increased. Note also that these patterns are reflected in the dependence of (*Ns*_*b*_)^*^ on *η* and *NU* as shown in Fig. 3, as well as the dependence of full *v* = 0 curves on *η* and *NU*, which can be seen in Fig. S2.

### E. Patterns of Molecular Evolution

Despite the fact that the *v* = 0 surface involves no change in the mean fitness of a population over time, the evolutionary dynamics of populations on the *v* = 0 surface can be far from neutral. These populations lie in a dynamic steady state involving accumulation of both beneficial and deleterious mutations, at rates which may differ substantially from the accumulation rate of neutral mutations. The fixation probabilities 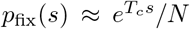, along with the MSSM approximation for determining *T*_*c*_, can be used to characterize the expected total rate F of (selected) *mutation fixation* of a population, both lying on or off of the *v* = 0 surface. In particular, with analogy to our characterization of a *v* = 0 surface, an *F* = *U* surface can be characterized on which selected mutations, on average, accumulate/fix as if they were entirely neutral (with faster-than-neutral accumulation of beneficial mutations precisely balancing slower-than-neutral accumulation of deleterious mutations). If we assume that synonymous mutations are neutral and nonsynonymous mutations are selected, the *F* = *U* surface can also be thought of as a *dN*/*dS* = 1 surface (which would typically be interpreted as evidence for neutral, or nearly neutral, evolution (Kryazhimskiy and Plotkin, 2008; Yang and Bielawski, 2000)).

The F = *U* surface is described by the equation 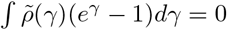, from which it follows that

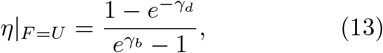

for the special case of a two-effect DFE. Note that η|_*F* =*U*_ increases monotonically with *γ*_*d*_, and decreases monotonically with *γ*_*b*_; thus, in contrast to *v* = 0 curves, *F* = *U* curves do not attain extrema (at least in the space of scaled effects). In contrast to the *v* = 0 surface, *γ*_*b*_ does not tend to 0 as *γ*_*d*_ → ∞, but instead tends to log (1 + 1/η). As a result, to obtain *F* = *U* curves in the space of unscaled effects, the “N_*e*_-based heuristic” described above, in which *T*_*c*_ is given by Eq. (11), does not apply; even at large *γ*_*d*_, the dynamics are not driven primarily by strong purifying selection on deleterious mutations. Instead, we can use *T*_*c*_ obtained with the MSSM approximation (that is, using Eq. (8)) over a larger range of Ns_*d*_ values. In Fig. 4, we plot a grid of fixation rates obtained from the same simulations (of populations subject to two-effect DFEs) considered in Fig. 2, along with corresponding *F* = *U* theory curves obtained using the MSSM approximation. Our theory curves quite accurately distinguish simulated parameter combinations in which *F* > *U* from those in which *F* < *U*. The same comparison is provided in Fig. S4, including populations with different N*U* values and populations subject to a two-exponential DFE. We note that, as compared to the *F* = *U* surface in the space of scaled effects, the behavior is slightly more complex in this case, and the F = *U* surface can be somewhat non-monotonic in certain cases (due to variation of *T*_*c*_ with Ns_*b*_ and Ns_*d*_).

**FIG. 4.**
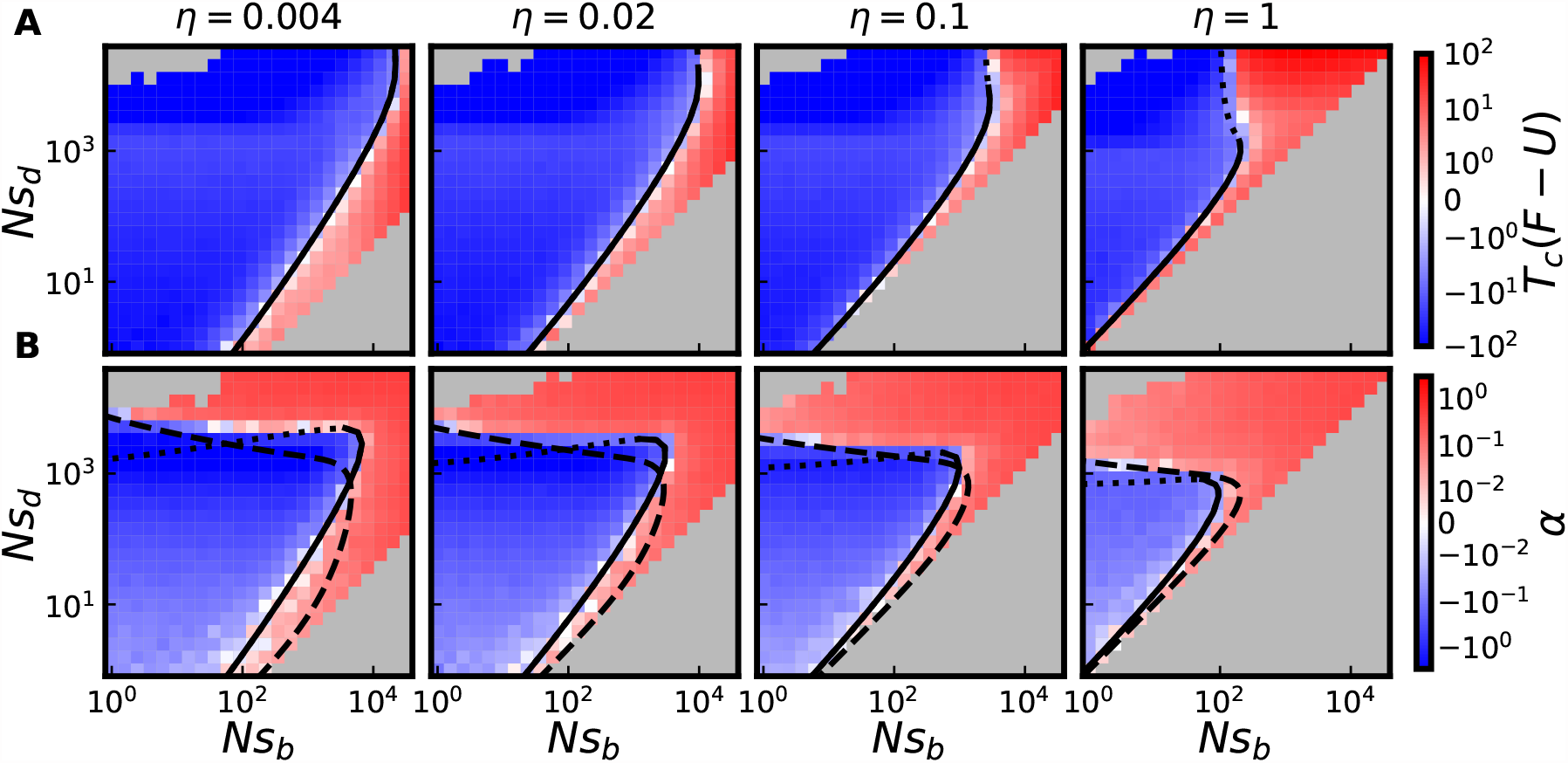
Cross sections of *F* = *U* and *α* = 0 surfaces. (A) Populations shown in Fig. 2A (which are subject to a two-effect DFE with *NU* = 10^4^) are colored by their values of *T*_*c*_(*F*− *U*) measured in simulations. (B) Populations shown in Fig. 2A are colored by measured values of *α* in simulations. In A and B, solid lines depict predictions of the MSSM approximation for the *F* = *U* and *α* = 0 surface, respectively, within its regime of validity; dotted lines denote MSSM predictions beyond its regime of validity. In B, dashed lines denote the corresponding predictions obtained using the *N*_*e*_-based heuristic described above.

Measurements of within-population genetic *diversity* enable another way of characterizing and potentially drawing inferences from a population. In particular, an imbalance in the number of nonsynonymous (and synonymous) polymorphisms observed, relative to the number of nonsynonymous (and synonymous) mutations fixed since divergence of two populations, is often interpreted as a signature of selection—either positive or negative—through the McDonald Kreitman test (McDonald and Kreitman, 1991). We can quantify a related imbalance through the statistic α, defined as

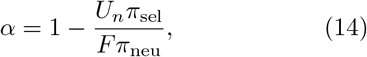

which resembles the McDonald-Kreitman statistic α_*MK*_, making an analogy between synonymous mutations and neutral mutations, and between nonsynonymous mutations and selected mutations. Here π_neu_ and π_sel_ denote levels of pairwise heterozygosity of neutral and selected mutations, respectively (with pairwise heterozygosity simply the average number of polymorphisms observed in a sample of two individuals). The statistic α_*MK*_ has often been used to estimate the fraction of substitutions in a population which are adaptive (Eyre-Walker, 2006). However, neither α_*MK*_ nor the statistic α defined here need be positive; α_*MK*_ < 0 is often observed and interpreted as evidence that purifying selection plays a dominant role in the evolution of a population (Charlesworth and Eyre-Walker, 2008). The α = 0 surface is thus a third surface on which the dynamics are at least ostensibly neutral in some capacity, and which can also be described using the MSSM approximation. We reproduce results of the MSSM approximation for π_neu_ and π_sel_ in the *SI Appendix*. Using these results, the α = 0 surface follows as

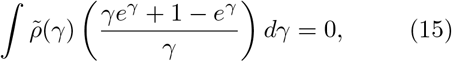

a consequence of which is that

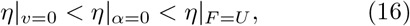

for any 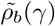 and 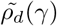. Thus, *v* > 0 and *F* < *U* on the α = 0 surface (and α < 0 and *F* < *U* on the *v* = 0 surface). Fig. 4 includes a comparison between predictions for the α = 0 surface—obtained using Eq. (15), along with Eq. (8) or Eq. (11)—and the values of α obtained in simulations, for populations subject to two-effect DFEs. To obtain predictions for the α = 0 surface in the space of unscaled effects, the “N_*e*_-based heuristic” is again useful for large Ns_*d*_ and the MSSM approximation (i.e. Eq. (8)) is useful for small Ns_*d*_ in relating *T*_*c*_ to N, although the specific patching we have employed in Eq. (12) to obtain the *v* = 0 surface does not quite carry over. We provide a comparison between α = 0 predictions and simulated α values for populations subject to a two-exponential DFE in Fig. S5; except for cases when *η* is small and Ns_*d*_ is large (and the MSSM approximation breaks down) agreement is qualitatively similar.

## IV. DISCUSSION

Previous efforts to treat the dynamics of both beneficial and deleterious mutations have largely done so by treating the two types of mutations in fundamentally different ways (Bachtrog and Gordo, 2004; Good and Desai, 2014; Goyal et al., 2012; Johnson and Barton, 2002; McFarland et al., 2013). These efforts typically start by making one of several assumptions about whether and how often deleterious mutations can fix, and how they impact they fixation probabilities of beneficial mutations. We lack an adequate understanding of the interplay between the two types of mutation in general, or even of which type of mutation will be more important in shaping the fitness trajectory of a population, given a particular distribution of fitness effects. Here, we have used our recently developed MSSM approximation, along with a simple *N*_*e*_-based heuristic, to characterize this balance between beneficial and deleterious mutations. Our description of the *v* = 0 surface applies under quite general conditions we have described above—essentially, as long as N*U*_*b*_ is sufficiently large, so that beneficial mutations enter the population sufficiently frequently. As we discuss in more detail below, the *v* = 0 surface is particularly relevant to the fate of a population at long evolutionary times: the *v* = 0 surface limits the ability of evolution to climb fitness landscapes—and thus, under certain conditions, determines the extent to which evolution acts as an optimization process.

We have found that the *v* = 0 constraint is concisely expressed in terms of the distribution of scaled fitness effects *T*_*c*_s available to a population. Expressed as such, the population size N is relevant only via its impact on the coalescence timescale *T*_*c*_. Alternatively, given a fixed *U* and *ρ*(*s*), a particular N_0_ can be identified such that *v* > 0 for *N* > *N*_0_ and *v* < 0 for *N* < *N*_0_. This has long been recognized in the context of *mutational meltdown* models (Lynch et al., 1993), which typically assume that decreases in fitness imply decreases in a population’s size, and thus further decreases in its fitness (or the opposite increase in its size, if the population instead increases in fitness initially). Our analysis immediately yields a critical *effective* population size *N*_*e*_ at which *v* = 0, by rearranging Eq. (5), for example; a similar critical effective population size is identified by Whitlock (2000). Because the MSSM approximation provides a relation between *T*_*c*_ and N, however, our analysis can also yield a critical *census* population size *N*_0_ at which v = 0, for a given *Uρ*(*s*).

Previous work has considered the balance between accumulation of beneficial mutations and deleterious mutations under more limiting assumptions (Goyal et al., 2012; Held et al., 2019; McFarland et al., 2014; Rice et al., 2015). Notably, Goyal et al. (2012) compute the fraction *U*_*b*_/(*U*_*b*_ + *U*_*d*_) of beneficial mutations to total mutations at which *v* = 0, given a single effect size s_*b*_ = s_*d*_ = s of both beneficial mutations and deleterious mutations. Goyal et al. (2012) also provide bounding arguments for the case in which the effect sizes of beneficial mutations and deleterious mutations differ, and briefly discuss the case in which distributions of fitness effects are more broad. The key idea of this and other single-effect approaches is that the single effect s that is modeled must be chosen as the most-likely (or in a sense, typical) effect size of a *fixed* mutation (Fisher, 2013; Good et al., 2012). Thus, use of a single-s approach to obtain a *v* = 0 constraint requires that the most-likely effect size of a fixed beneficial mutation at least roughly matches the most-likely effect size of a fixed deleterious mutation.

Rice et al. (2015) have found that, under the assumption of no epistasis (i.e., assuming a genome of finite size where mutations at the different loci do not interact epistatically) this is precisely to be expected after long evolutionary timescales. Their basic argument is simple: if beneficial mutations of a particular effect size are more likely to fix than deleterious mutations of that effect size, those mutational opportunities will be depleted faster (or vice versa). This will continue until the distribution of fixed beneficial effects precisely matches the distribution of fixed deleterious effects, at which point adaptation will come to a halt (*v* = 0) and the DFE can be described as “evolutionarily stable”. Therefore, a single-s approach is perhaps appropriate in describing the approach to an evolutionary attractor at long times resulting from a population running out of beneficial mutations. In the presence of epistasis, however, a mutation not only enables a back mutation of the opposite effect, but can also alter the full distribution of fitness effects available to an individual. As a result, the distributions *ρ*_*b*_(s) and *ρ*_*d*_(s) can change in independent ways over the course of evolution. Our analysis—which makes no assumption that *ρ*_*b*_(s) and *ρ*_*d*_(s) are similar in scale—is thus more applicable to a description of the *v* = 0 constraint in the presence of widespread epistasis. In particular, as we now discuss, our analysis provides a framework to work out the implications of various proposed patterns of *fitness-mediated epistasis* (Good and Desai, 2015), in which *ρ*(s) varies systematically with the fitness of a population.

To do so requires an additional assumption of how the parameters of a population depend on fitness.

For instance, in the presence of *diminishing-returns* epistasis (in which beneficial effects become systematically *weaker* as the fitness of a population increases) populations are constrained to lie on a horizontal line in the space *Ns*_*d*_ vs. *Ns*_*b*_ depicted in Fig. 2 and Fig. 4. If a population starts out in the *v* > 0 region, its mean fitness will increase and its value of Ns_*b*_ will subsequently decrease, until the population converges to the *v* = 0 surface. Qualitatively, then, the implications of diminishing-returns epistasis are similar to the implications of a declining fraction of beneficial mutations (which are considered by Goyal et al. (2012)): as evolution proceeds, the population will approach the *v* = 0 surface (either from the *v* < 0 region or the *v* > 0 region); the *v* = 0 surface is thus a stable evolutionary attractor. On the other hand, a pattern of *increasing-costs* epistasis (in which deleterious fitness effects systematically increase with fitness) corresponds to a vertical line in the space Ns_*d*_ vs. Ns_*b*_. The behavior in this case is more complex, depending on the location of the starting point in relation to the *v* = 0 ridgeline: if 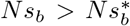,a long term evolutionary attractor does not exist, while if 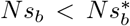, two fixed points exist, one of which is stable and one of which is unstable. One could also imagine a pattern of *decreasing-costs* epistasis, in which deleterious fitness effects become systematically *smaller* as a population increases in fitness (e.g. if deleterious mutations are thought of as reversions of beneficial mutations for which a pattern of diminishing-returns epistasis exists). The behavior in this case is similar to that of increasing-costs epistasis, but the stablity (and instability) of the two fixed points is swapped.

It is not yet entirely clear which, if any, of these simple patterns best describe the dominant patterns of epistasis in natural populations. More generally, a curve through the parameter space—parameterized by fitness—can be assumed, on which populations are constrained to lie. The intersection(s) of these curves with the *v* = 0 surface determine the long-term evolutionary fixed points, the stability of which can be determined straightforwardly. In principle, these curves may covary along several dimensions (e.g. with quantities such as the overall mutation rate, the relative fraction of beneficial to total mutations, and other quantities such as the shapes of beneficial and deleterious DFEs). In Fig. 5, as a schematic we illustrate the consequences of a relatively simple pattern involving both diminishing-returns epistasis and increasing-costs epistasis; as recently argued by Lyons et al. (2020) and Reddy and Desai (2021), these two trends both emerge from a simple null model of pervasive microscopic epistasis. Populations are simulated starting from a range of five initial conditions in the parameter space; in each case, s_*b*_ decreases with the fitness of an individual and s_*d*_ increases with fitness of an individual, with the product s_*b*_s_*d*_ assumed constant (see *Materials and Methods* for details). Note that for two of the five initial conditions, the population would never approach the *v* = 0 surface (but its rate of adaptation does slow down over time). For the remaining three initial conditions, the population approaches the *v* = 0 surface, either from higher fitness or from lower fitness, and in particular approaches the 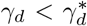 portion of the *v* = 0 surface described by the MSSM approximation.

**FIG. 5.**
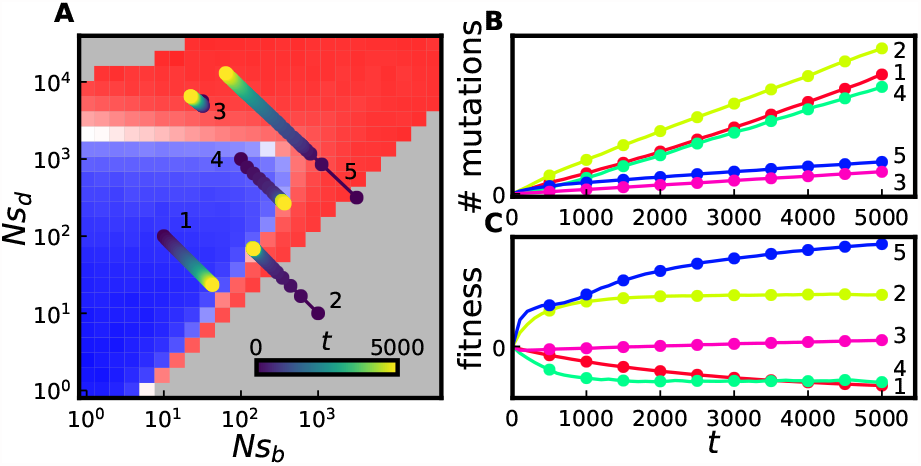
(A) Simulated trajectories of evolving populations subject to both diminishing-returns and increasing-costs epistasis, overlaid on panel C of Fig. 2. Populations are subject to two-effect DFEs with *NU* = 10^4^ and *η* = 0.1. In all cases, the product (*Ns*_*b*_)(*Ns*_*d*_) is assumed constant as a population evolves; depending on a population’s initial values of *Ns*_*b*_ and *Ns*_*d*_ it may or may not approach the *v* = 0 surface at long times. The color of a filled circle denotes the time (in generations) a population had a given set of parameters. (B) Mutation accumulation and (C) fitness trajectories of the simulated populations depicted in A. Despite a declining rate of fitness change observed in all cases, rates of (selected) mutation accumulation remain roughly constant.

In all cases shown in Fig. 5, selected mutation accumulation proceeds throughout at a roughly constant rate, although, consistent with fixation rates measured in simulations and shown in Fig. 4, those populations which approach the *v* = 0 surface have a higher long-term steady state fixation rate of new mutations. Those populations which approach the *v* = 0 surface can be thought to undergo rapid molecular evolution at steady state, in that F ∼ *U*_*b*_ + *U*_*d*_ is possible. In contrast, those populations which do not approach the *v* = 0 surface can instead end up with a much smaller fixation rate F ∼ *U*_*b*_ (with F = *U*_*b*_ if deleterious mutations are strong enough that all are purged by selection, and beneficial mutations weak enough that they accumulate entirely neutrally). Broadly speaking, these patterns—a rate of fitness increase which declines over time (Couce and Tenaillon, 2015), and in particular, the maintenance of a roughly constant rate of mutation accumulation despite a declining rate of fitness increase (Barrick et al., 2009; Good et al., 2017; Kryazhimskiy et al., 2014)—have been observed in multiple microbial evolution experiments. Our analysis provides a way to identify regions of the parameter space in which these and similar observed patterns are possible, or alternatively, to yield constraints on the dominant modes of fitness-mediated epistasis given an observed fitness and/or mutation accumulation trajectory.

Using arguments along the lines described above, our analysis can be used to predict the flow of a population through parameter space, given a complete characterization of any form of fitness-mediated epistasis—that is, of *ρ*(s|X). Crucially, we make the assumption of *slow epistasis*, such that *ρ*(s|X) can be treated as uniform within the population, and constant in time, in identifying its corresponding rate of fitness increase. This assumption means that we neglect the possibility that individual mutations could lead to specific shifts in *Uρ*(*s*), which could then themselves be subject to selection. For example, a lineage could arise that has access to more (or stronger-effect) beneficial (or deleterious) mutations than other individuals within the population, and this lineage could then be subject to second-order selection. Analyzing this effect is an interesting topic for other work, and has recently been addressed by Ferrare and Good (2023) using a related theoretical framework.

## V. MATERIALS AND METHODS

To validate our predictions, we conducted individual-based Wright-Fisher simulations. Simulations were performed using code available at https://github.com/mjmel/mssm-sim and used by Melissa et al. (2022). Simulations consist of a mutation step and a reproduction step repeated each generation. In the mutation step, each individual acquires a Poisson-distributed number of mutations with mean *U*; the effect of each mutation is independently drawn from the distribution *ρ*(s) and increments (or decrements) an indiviual’s log-fitness *X*. Purely neutral mutations are also introduced at rate *U*_*n*_. The identities and fitness effects of the mutations carried by each individual are tracked. In the reproduction step, individuals are resampled with replacement with probabilities proportional to their fitnesses *e*^*X*^. Populations are initialized clonally and the number of generations which elapses before the first fixation of a mutation is recorded, setting the *epoch length*. Simulations are run for up to 100 epochs, with the mean fitness and heterozygosities—both of neutral mutations and of selected mutations—recorded at each epoch. Simulations which for which fewer than 10 epochs have been reached after a runtime of 24 hours are discarded from our analysis.

For a given value of *η* and N*U*, we simulated parameter combinations lying on a grid of Ns_*b*_ and Ns_*d*_ values (depicted in Fig. 2, for example). For each value of *η* and N*U*, we extracted the coordinates of its ridgeline point, both in the space of scaled effects and in the space of unscaled effects, for comparison with theory. To do so in the space of scaled effects, we take the point with the largest *γ*_*b*_ value such that *v* < 0; the *γ*_*b*_ and *γ*_*d*_ of this point are then recorded as 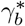 and 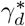, respectively. In the space of unscaled effects, we take (*Ns*_*b*_)^*^ as the largest value of *Ns*_*b*_ such that v < 0 for some parameter combination; (*Ns*_*d*_)^*^ is then taken as the median value of Ns_*d*_ for those parameter combinations such that *Ns*_*b*_ = (*Ns*_*b*_)^*^ and *v* < 0.

We also conducted simulations in which the available fitness effects *s*_*b*_ and s_*d*_ depend on fitness. These simulations are identical to those described above, except after the occurrence of each mutation, the effect magnitudes of the next available beneficial and deleterious mutations are updated accordingly. In all cases, we held the product s_*b*_s_*d*_ constant with *s*_*b*_ ∝ e^−*X/*5^, where *X* denotes an individual’s log-fitness. A similar functional dependence of *s*_*b*_ on *X* is found by Wiser et al. (2013) to describe *E.coli* populations in the LTEE experiment. These simulations were run for a total of 5000 generations, with measurements of fitness and the number of fixed mutations recorded every 100 generations.

## IV. ACKNOWLEDGEMENTS

We thank Benjamin Good and Ivana Cvijovic for many helpful comments and suggestions. MMD acknowledges support from NSF Grant PHY-1914916 and NIH grant R01GM104239. Simulations were conducted on the FASRC Cannon cluster supported by the FAS Division of Science Research Computing Group at Harvard University.

## SI Appendix

### I. REVIEW OF THE MODERATE SELECTION, STRONG MUTATION APPROXIMATION

In this section we provide a brief overview of the moderate selection, strong mutation (MSSM) approximation, which can be used to analyze evolutionary dynamics in the MSSM regime. This summarizes the key results of Melissa et al. (2022) that are relevant to our characterization of the *υ* = 0 constraint.

A central aim is to predict, given the parameters *N* and *Uρ*(*s*), the dynamical quantities *υ* and *p*_fix_(s), along with the timescale *T*_*c*_ of pairwise coalescence. A key result of Melissa et al. (2022) is that under the MSSM approximation,

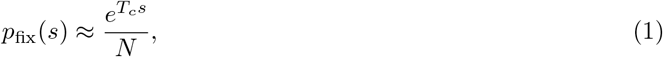

for *s* ≪ *b*, where *T*_*c*_ is defined by

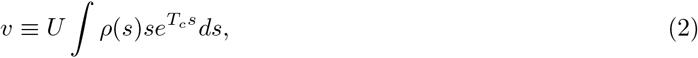

and *b* is defined by

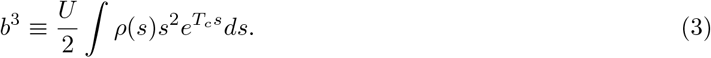

Note that to apply the MSSM approximation, *ρ*(s) must fall off exponentially, or faster than exponentially, with large positive s; otherwise, the above integrals do not converge and *T*_*c*_ is not well-defined. The quantity *T*_*c*_ is related to the underlying parameters *N* and *Uρ*(s) through the following system of equations for *T*_*c*_ and the interference threshold *x*_*c*_ (the fitness advantage above which individuals can fix largely unhindered by clonal interference, as well as, roughly speaking, the typical fitness advantage of the most-fit individuals in the population):

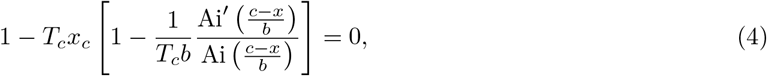

and

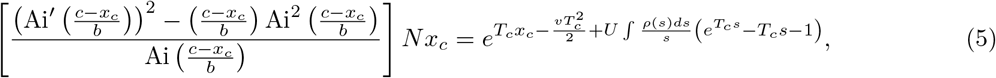

where

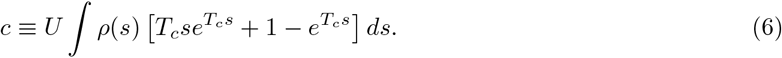

Eq. (4) and Eq. (5) are used throughout this work to obtain theory predictions under the MSSM approximation. While numerical solution of these equations is straightforward, it is sometimes useful to further approximate these two equations as

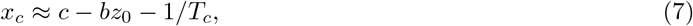

and

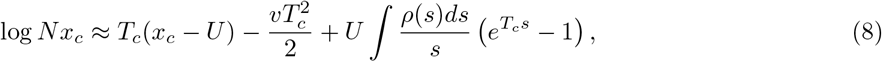

respectively, which holds in the regime of validity of the MSSM approximation (which we discuss below).

While defined by Eq. (2), the quantity *T*_*c*_ approximately corresponds to ⟨*T*_2_⟩ /2—one-half the average time since two randomly chosen individuals in a population share a common ancestor—and can be interpreted as a *coalescence timescale*. In particular, the pairwise neutral heterozygosity *π*_neu_ (the average number of neutral genetic differences among pairs of individuals in a population) evaluates approximately to 4*T*_*c*_*U*_*n*_; the relation ⟨*T*_2_⟩/2 ≈ *T*_*c*_ follows since *π*_neu_ = 2 ⟨*T*_2_ ⟩ *U*_*n*_. The pairwise *selected* heterozygosity (which counts *selected* genetic differences, occurring at rate *U*, with fitness effects drawn from *ρ*(s)) is given by

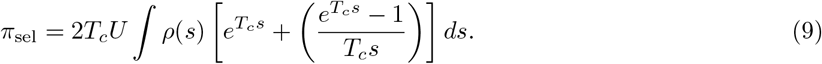

Results have also been obtained for the full site frequency spectrum (which gives the expected number of mutations present in a sample at a given frequency, and from which additional statistics of genetic diversity can be computed) for both neutral mutations and selected mutations. The fixation rate of neutral mutations is simply given by *U*_*n*_, while the fixation rate of selected mutations is given by 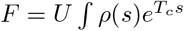; from these prediction, a prediction for *α* (as defined in the main text) follows.

Approximate validity of the MSSM approximation requires that *T*_*c*_b ≫ 1 and that

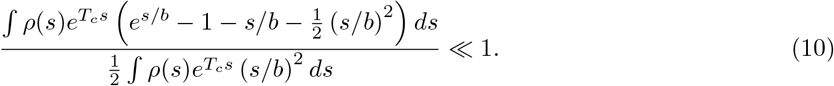

For populations subject purely to beneficial mutaitons, or purely to deleterious mutations, the second condition is roughly encapsulated by the condition 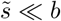, where 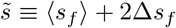 denotes the largest “typical” effect size of a fixed mutation, as well as by the requirement that *ρ*(s) falls off exponentially, or faster than exponentially with large positive *s*. The quantity ⟨*s*_*f*_ ⟩ denotes the *average* effect size of a fixed mutation, and the quantity Δ*s*_*f*_ denotes the *standard deviation* in effect sizes of fixed mutation. When beneficial and deleterious mutations both occur in a population, separate largest “typical” effects 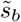 and 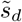 can defined for beneficial and deleterious mutations, and both 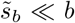 and 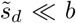 must be satisfied. These conditions of validity must be satisfied self-consistently; provided that these conditions are met, the distribution *ρ*_*f*_ (s) of *fixed* mutational effects follows as 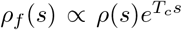 within the “bulk” of *ρ*_*f*_ (s)—the region dominating ∫*ρ*_*f*_ (s)ds. These conditions ensure that the distribution of relative fitnesses f(x) is well-approximated by

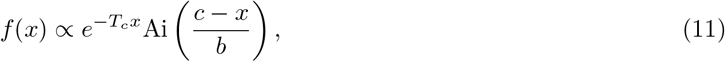

and that the fixation probability *w*(*x*) of an individual with relative fitness *x* is well-approximated by

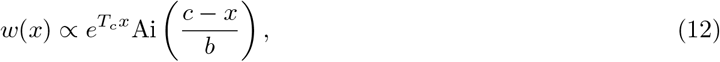

within the important region of fitness space (the “fixation class”, near the “nose” of the fitness distribution) which produces the majority of future common ancestors of the population. This region of fitness space (which dominates the integral ∫ *f*(*x*)*w*(*x*)d*x*) is particularly important because Eq. (8) is obtained by enforcing that 1/*N* =∫ *f*(*x*)*w*(*x*)d*x* (i.e., that on average, one individual in the population at any given time will eventually fix).

### II. APPLICABILITY OF THE MSSM APPROXIMATION ALONG THE *υ* = 0 RIDGELINE

In the main text, we argued that the *υ* = 0 ridgeline (in the space of scaled fitness effects) is given by the curve 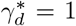 and 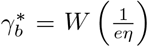, parameterized by *η*. Here, we evaluate the quantities *T*_*c*_*b*, s_*d*_/b and *s*_*b*_/*b* along this *υ* = 0 ridgeline, which we then use to comment on the validity of the MSSM approximation in describing the *υ* = 0 ridgeline.

From the definition of b in Eq. (3), it follows that

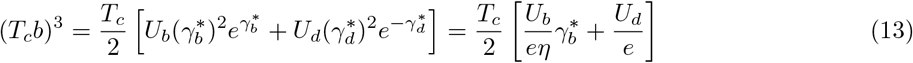

on the *υ* = 0 ridgeline, which in turn simplifies to

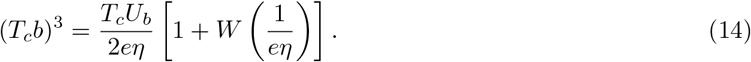

As a consequence, *T*_*c*_*b* ≫ 1 and *s*_*d*_/*b* ≪ 1 if *T*_*c*_*U*_*b*_ ≫ 1 (assuming *η* < 1). Furthermore, given Eq. (14) we have

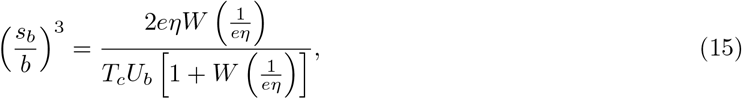

with the right-hand side of Eq. (15) bounded above by 0.4/(*T*_*c*_*U*_*b*_); thus, *s*_*b*_/*b* ≪ 1 along the *υ* = 0 ridgeline if T_*c*_*U*_*b*_ ≫ 1.

### III. DISTRIBUTIONS OF FITNESS EFFECTS

Here, we consider simple classes of distributions of fitness effects, simplifying expressions obtained in the main text, and justifying the use of the MSSM approximation in describing the *υ* = 0 ridgeline. We focus our attention on the case in which effect sizes of both beneficial mutations and deleterious mutations are drawn from gamma distributions, potentially differing in shape and scale. That is, we assume

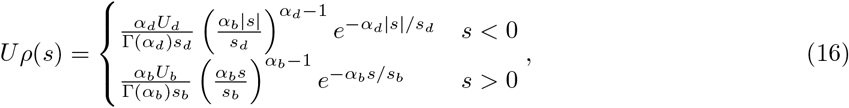

with *s*_*b*_ and *s*_*d*_ the mean available effect size of a beneficial mutation, or a deleterious mutation, respectively— that is, *s*_*b*_ ≡ ∫*ρ*_*b*_(*s*)*sds* and *s*_*d*_ ≡ ∫ *ρ*_*d*_(*s*)|*s*|*ds*. The parameters *α*_*b*_ and *α*_*d*_ comprise the shape parameters of the respective gamma distributions; the choice *α*_*b*_ = 1 (or *α*_*d*_ = 1) corresponds to an exponential distribution of beneficial (or deleterious) fitness effects; the limit *α*_*b*_ → ∞ (or *α*_*d*_ → ∞) corresponds to a delta distribution of beneficial (or deleterious) fitness effects.

Given the *Uρ*(s) specified in Eq. (16), the scaled rate of adaptation T_*c*_*υ* follows as

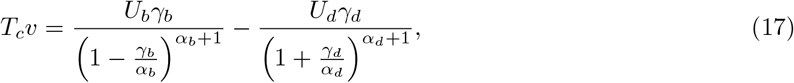

where the quantities *γ*_*b*_ and *γ*_*d*_ denote the average scaled effect sizes of beneficial mutations, and of deleterious mutations, respectively. Note that the requirement *γ*_*b*_ < *α*_*b*_ emerges, a consequence of Eq. (2), which defines *T*_*c*_. The *υ* = 0 surface then follows as

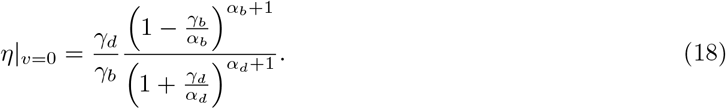

The fixation rate *F* can also be computed, using Eq. (1), with the result

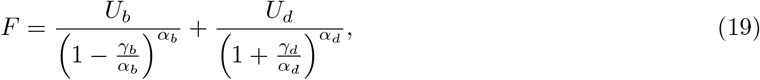

so that

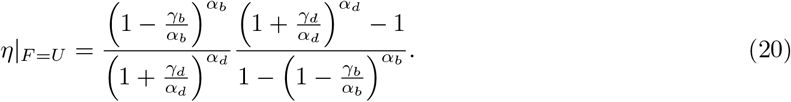

Finally, the quantity (*T*_*c*_*b*)^3^ simplifies to

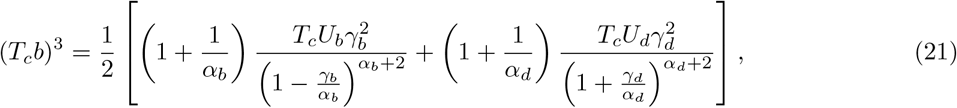

where in Eq. (21), we do not impose the constraint that *υ* = 0. Below, we focus our attention on the case in which both beneficial mutations and deleterious mutations are drawn from exponential DFEs (so that *α*_*b*_ = *α*_*d*_ = 1).

#### A. Exponentially-distributed fitness effects

In the case *α*_*b*_ = *α*_*d*_ = 1, the *υ* = 0 surface (in the space spanned by axes *η, γ*_*b*_ and *γ*_*d*_) is described by

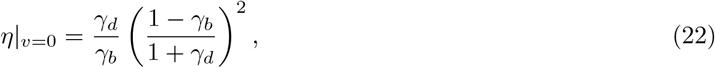

and the *υ* = 0 ridgeline is given by

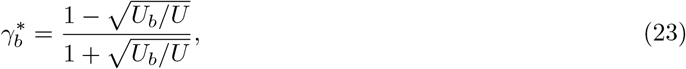

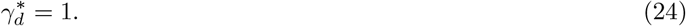

Because 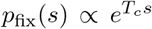 (at least for the majority of fixed mutations, assuming validity of the MSSM approximation), the distribution of *fixed* beneficial effects is also an exponential distribution, to a good approximation (and likewise for deleterious mutations); we have ⟨*s*_*f*_ ⟩_*b*_ = (Δ*s*_*f*_)_*b*_= *s*_*b*_/(1 − *T*_*c*_s_*b*_) and ⟨*s*_*f*_ ⟩_*d*_ = (Δ*s*_*f*_)_*d*_ = *s*_*d*_/(1 + *T*_*c*_*s*_*d*_). At the extremal point, we thus have

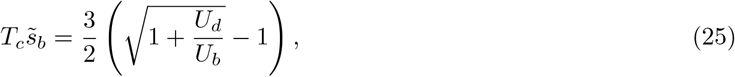

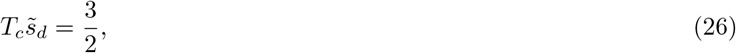

and

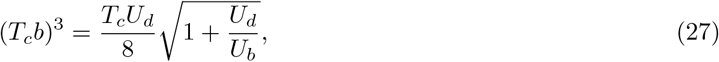

where 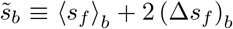 and 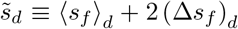. From Eq. (25) and Eq. (27) it follows that at the extremal point, 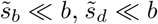 and *T*_*c*_*b* ≫ 1—and thus the conditions of validity of the MSSM approximation are satisfied—if *T*_*c*_*U*_*b*_ ≫ 1 (under the additional assumption that *U*_*d*_ ≥ *U*_*b*_).

We note that in the case of exponential DFEs, Eq. (20) simplifies to

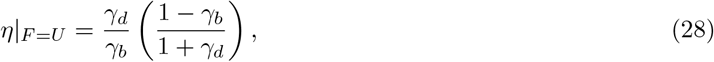

and *η*|_*α*=0_ can be computed using Eq. (9), with the result

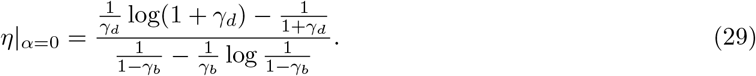

#### B. Arbitrary DFE

More generally, we can consider arbitrary DFEs *ρ*_*b*_(*s*) and *ρ*_*d*_(*s*) of beneficial and deleterious mutations. If we assume these DFEs have fixed shapes and consider changes in their scale, we can visualize the *υ* = 0 constraint as a 2-dimensional surface within a 3-dimensional parameter space parameterized by *η, γ*_*b*_ and *γ*_*d*_ (where here *γ*_*b*_ denotes the average scaled effect size of a beneficial mutation, and likewise for *γ*_*d*_). As in the case of single effects, a larger *γ*_*b*_ implies a more positive rate of adaptation, for fixed *η* and *γ*_*d*_. Likewise, an intermediate scale of deleterious effects is maximally impactful. For a particular shape of deleterious DFE, the maximally impactful deleterious (scaled) DFE 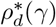, which lies on a *υ* = 0 ridgeline, satisfies

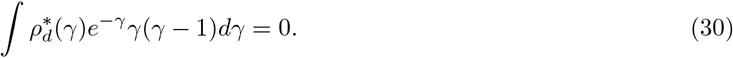

Because e^−*γ*^/*N* gives the fixation probability of a deleterious mutation with scaled effect *γ*, Eq. (30) has a simple interpretation in terms of the deleterious scaled effects which fix: on the *υ* = 0 ridgeline, the *mean-squared* fixed scaled deleterious effect equates to the *mean* fixed scaled deleterious effect. This further implies that 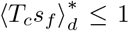 on the *υ* = 0 ridgeline (where s_*f*_ denotes a fixed mutational effect, and the expectation value averages over all deleterious fixed effects); note that this can be seen by ensuring positivity of the *variance* in fixed scaled deleterious effects, which equals 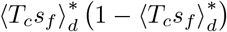, given Eq. (30). Thus, the average fixed effect of the maximally impactful deleterious DFE, 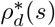, is subject only to moderate or weak selection. The upper bound 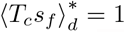 is achieved for the two-effect DFE considered above, while for the two-exponential DFE we have 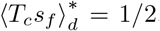. For the corresponding beneficial DFE, 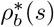, lying on the *υ* = 0 surface, a bound on 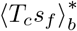 is less easily established without specifying the shape of *ρ*_*b*_(*s*).

### IV. EVOLUTIONARILY STABLE DFE

In the main text, we identified constraints on *Uρ*(*s*) (or perhaps on *Uρ*(*γ*), the distribution of scaled fitness effects) that yield *υ* = 0 and analyzed the resulting *υ* = 0 surface. In this fashion, *ρ*_*b*_(s) and *ρ*_*d*_(s) are treated as parameters which could in principle be varied independently. In some of the simplest models of genome evolution, however, *ρ*_*b*_(*s*) and *ρ*_*d*_(*s*) are not independent. For example, assuming a genome of finite length *L* with no epistasis, the fixation of a beneficial mutation creates an opportunity for a deleterious back-mutation of the same magnitude, and vice versa. Rice et al. (2015) consider the resulting “evolutionarily stable” DFE that is reached at long times once, for each magnitude of fitness effect |s|, beneficial and deleterious mutations reach a state of detailed balance, in which *ρ*_*b*_(*s*)p_fix_(*s*) = *ρ*_*d*_(*s*)*p*_fix_(*s*) (and consequently, the distribution of *fixed* beneficial mutations matches that of fixed deleterious mutations). A consequence of this detailed balance is that *υ* = 0 for the evolutionarily stable DFE; here we discuss the application of the MSSM approximation to this state.

Given the underlying distribution *ρ*_0_(|*s*|) of absolute effects |*s*|, 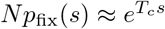 implies that

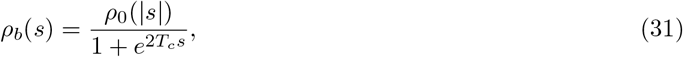

and

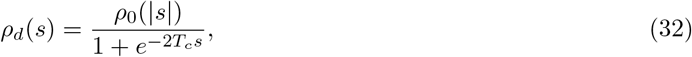

so that 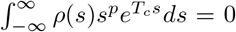 for odd *p*. The average effect size ⟨*s*_*f*_ ⟩ of a fixed beneficial mutation (which matches the average effect size of a fixed deleterious mutation) evaluates to

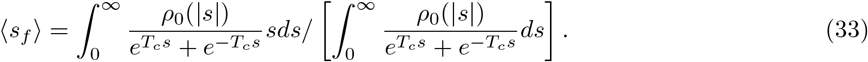

Below, we evaluate ⟨*s*_*f*_ ⟩ and related expressions for the class of stretched exponential DFEs 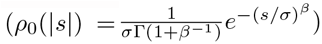 in the *T*_*c*_*σ*→ ∞ limit. The quantity ⟨ *s*_*f*_ ⟩ simplifies to

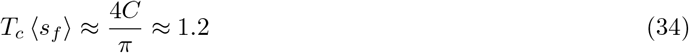

where *C* denotes Catalan’s constant. The mean-squared fixed effect (of beneficial mutations, and also of deleterious mutations) is given by

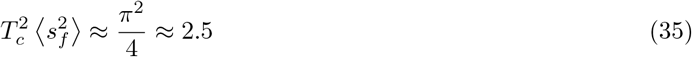

from which it follows that *T*_*c*_Δ*s*_*f*_ ≈ 1.1. In the same limit,

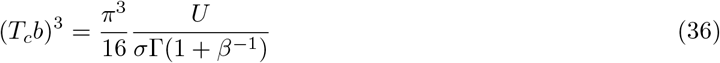

and so 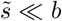 and *T*_*c*_*b* ≫ 1 in the *a* → ∞ limit if *U* ≫ σ.

---

**FIG. S1.**
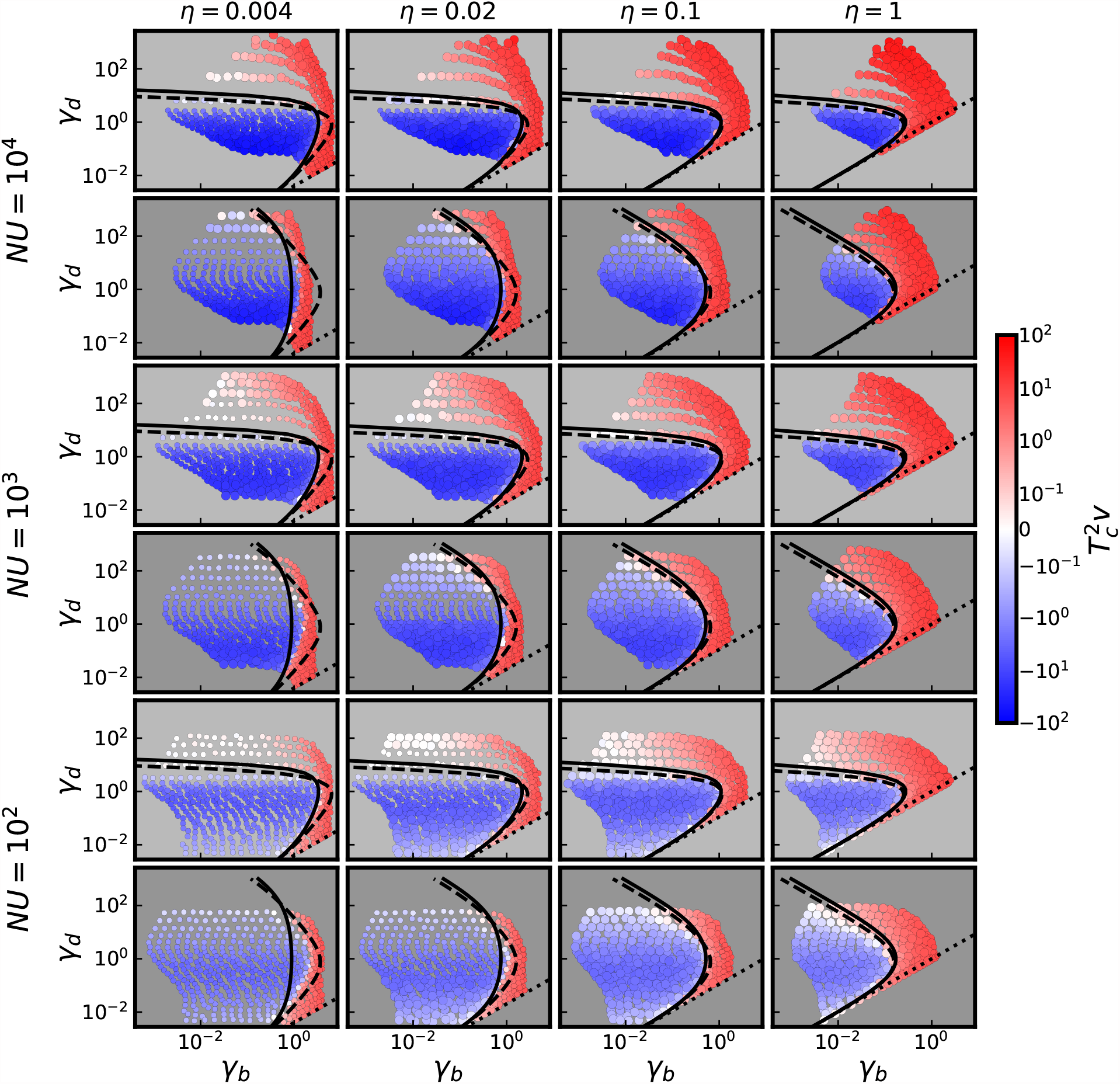
Cross sections of the *υ* = 0 surface in the space of scaled fitness effects. Populations are colored according to their values of 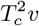 observed in simulations. For panels colored with a dark grey background (first, third and fifth rows) populations are subject to a two-effect DFE; for panels colored with a light grey background (second, fourth and sixth rows) populations are subject to a two-exponential DFE. The parameters *NU* and *η* for simulated populations in a given panel are as denoted on the left-hand side and top of the figure, respectively. Populations with *T*_*c*_*U*_*b*_ *<* 1*/*2 (which, at the *υ* = 0 ridgeline, suggests the MSSM approximation may break down) are denoted by points of smaller size. Solid curves denote predictions of the MSSM approximation, dashed lines denote predictions obtained using the standard formula for fixation probabilities assuming independently evolving loci, and dotted lines are the lines *γ*_*d*_ = *ηγ*_*b*_.

**FIG. S2.**
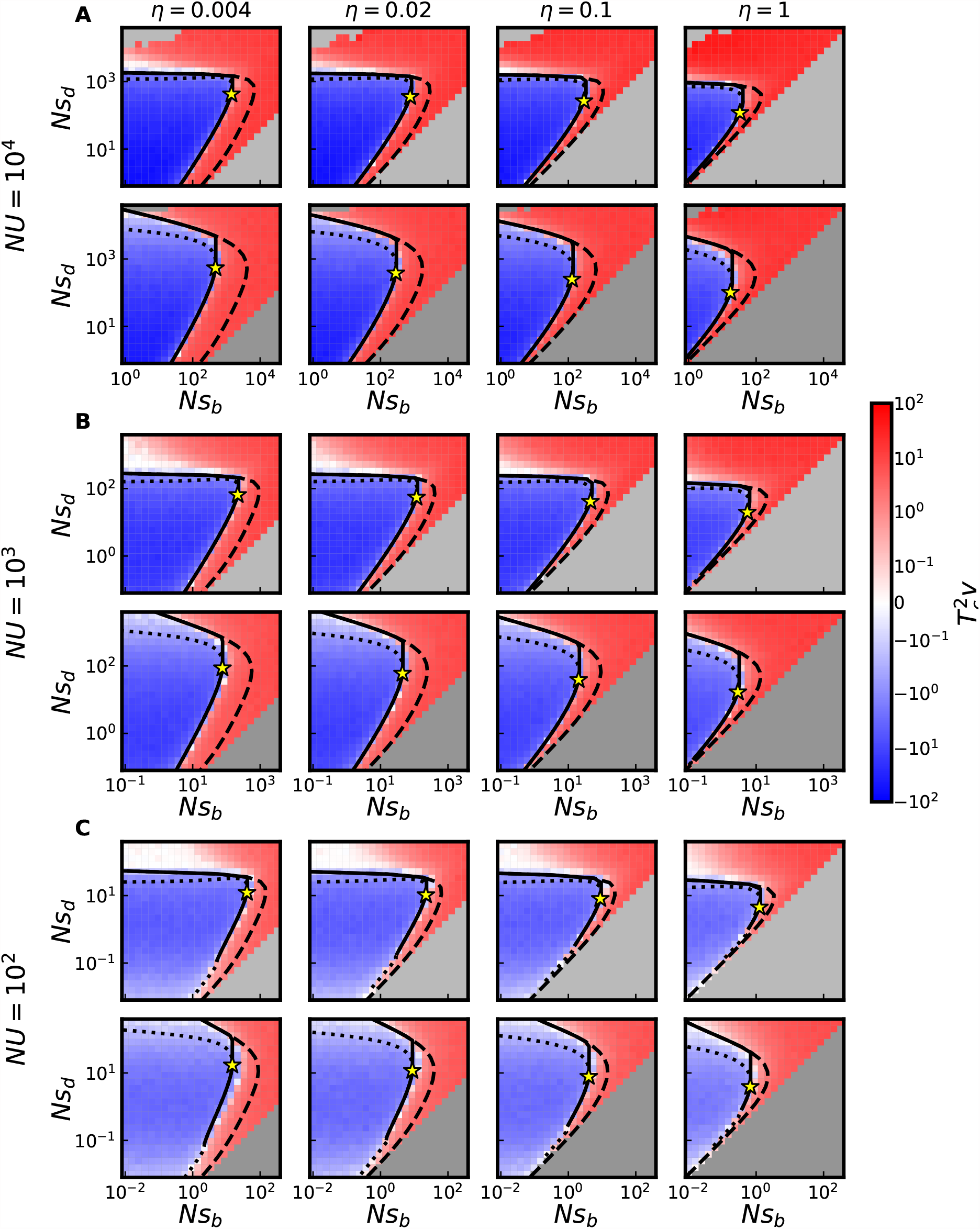
Cross sections of *υ* = 0 surface as obtained in simulations. Simulated populations have parameters *η* as depicted at the top of a given column, and *NU* = 10^4^ (in A), *NU* = 10^3^ (in B), or *NU* = 10^2^ (in C). Populations in the first, third and fifth rows (light grey background) are subject to a two-effect DFE; populations in the second, fourth and sixth rows (dark grey background) are subject to a two-exponential DFE. Solid curves denote piecewise-defined *υ* = 0 predictions given in the main text. Stars denote points at which MSSM approximation predicts *υ* = 0 and 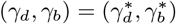. Dashed lines are predictions obtained using our “*N*_*e*_-based heuristic”. Dotted lines are MSSM predictions of *v* = 0 curves obtained for parameters such that *T*_*c*_*b <* 1 or *Ns*_*d*_ *>* (*Ns*_*d*_)^∗^.

**FIG. S3.**
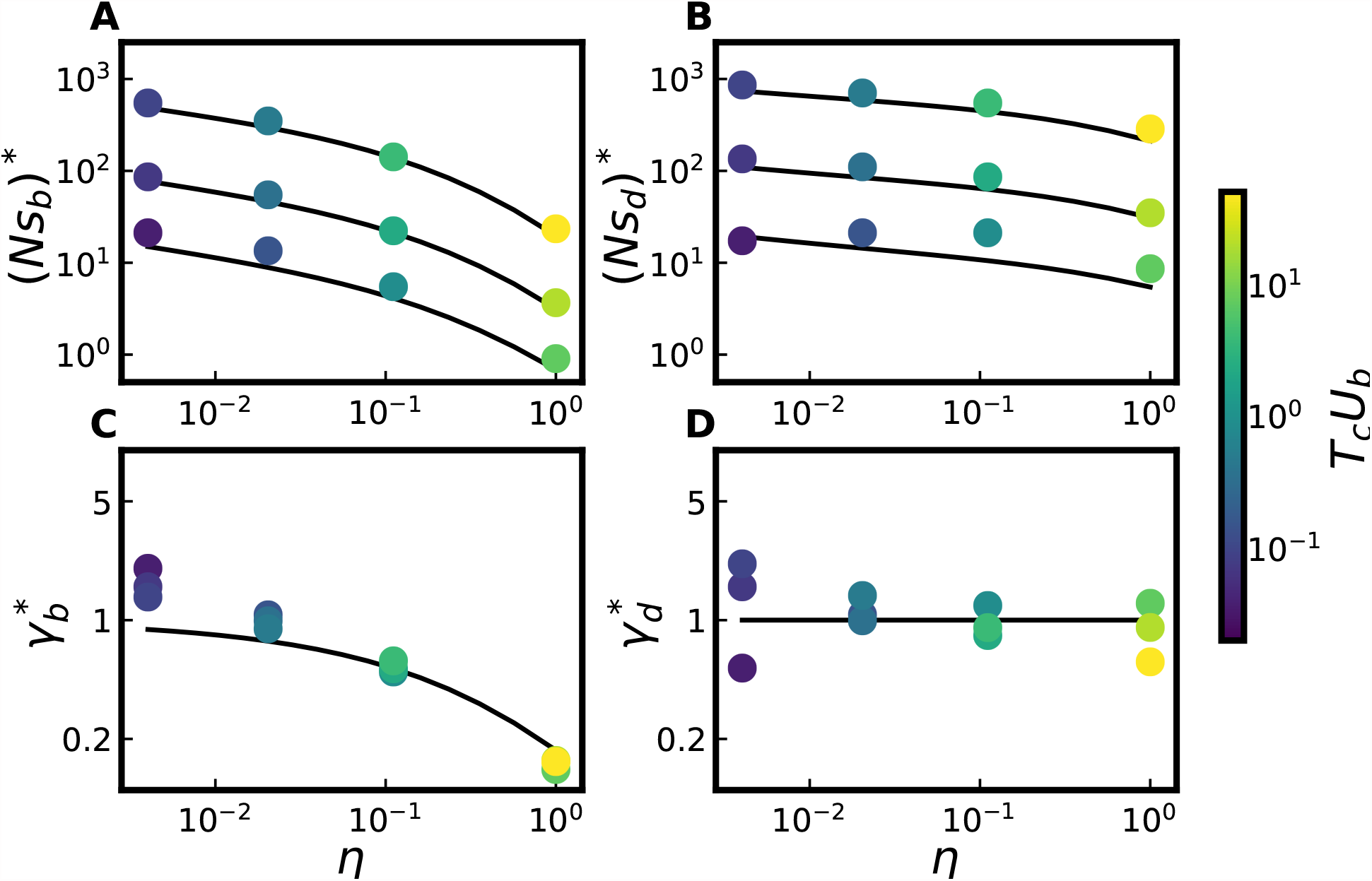
Comparison between simulations and MSSM predictions for, in panels A and B, the extremal point ((*Ns*_*b*_)^∗^, (*Ns*_*d*_)^∗^) and, in panels C and D, the ridgeline point 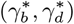. Each point corresponds to a panel of Fig. S2 consisting of populations subject to a two-exponential DFE, for particular values of *η* and *NU*. In both A and B, the three solid curves are theory curves for *NU* ∈ {10^2^, 10^3^, 10^4^} (with the top curve corresponding to *NU* = 10^4^ and the bottom curve corresponding to *NU* = 10^2^). Each point is colored according to its value of *T*_*c*_*U*_*b*_ at 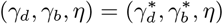.

**FIG. S4.**
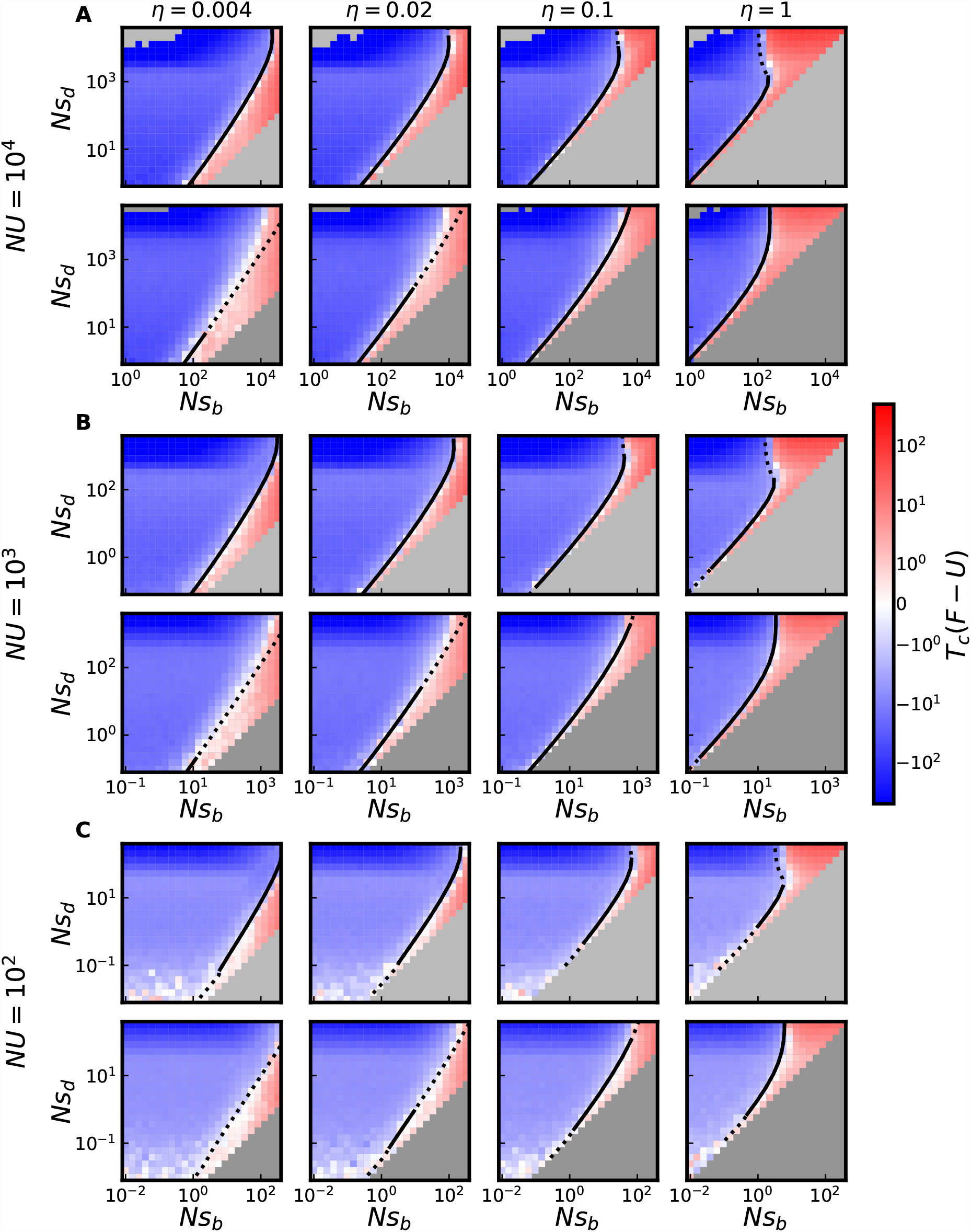
Cross sections of *F* = *U* surface as observed in simulations. Points are colored by values of *T*_*c*_(*F* − *U*) as observed in simulations. Values of *NU* for populations displayed in *A, B* and *C* are denoted on the left-hand side. Populations in the first, third, and fifth row are subject to two-effect DFEs; populations in the second, fourth, and sixth rows are subject to two-exponential DFEs. Solid curves denote MSSM predictions for *F* = *U* surface, with dotted lines denoting these predictions beyond the regime of validity of the MSSM approximation.

**FIG. S5.**
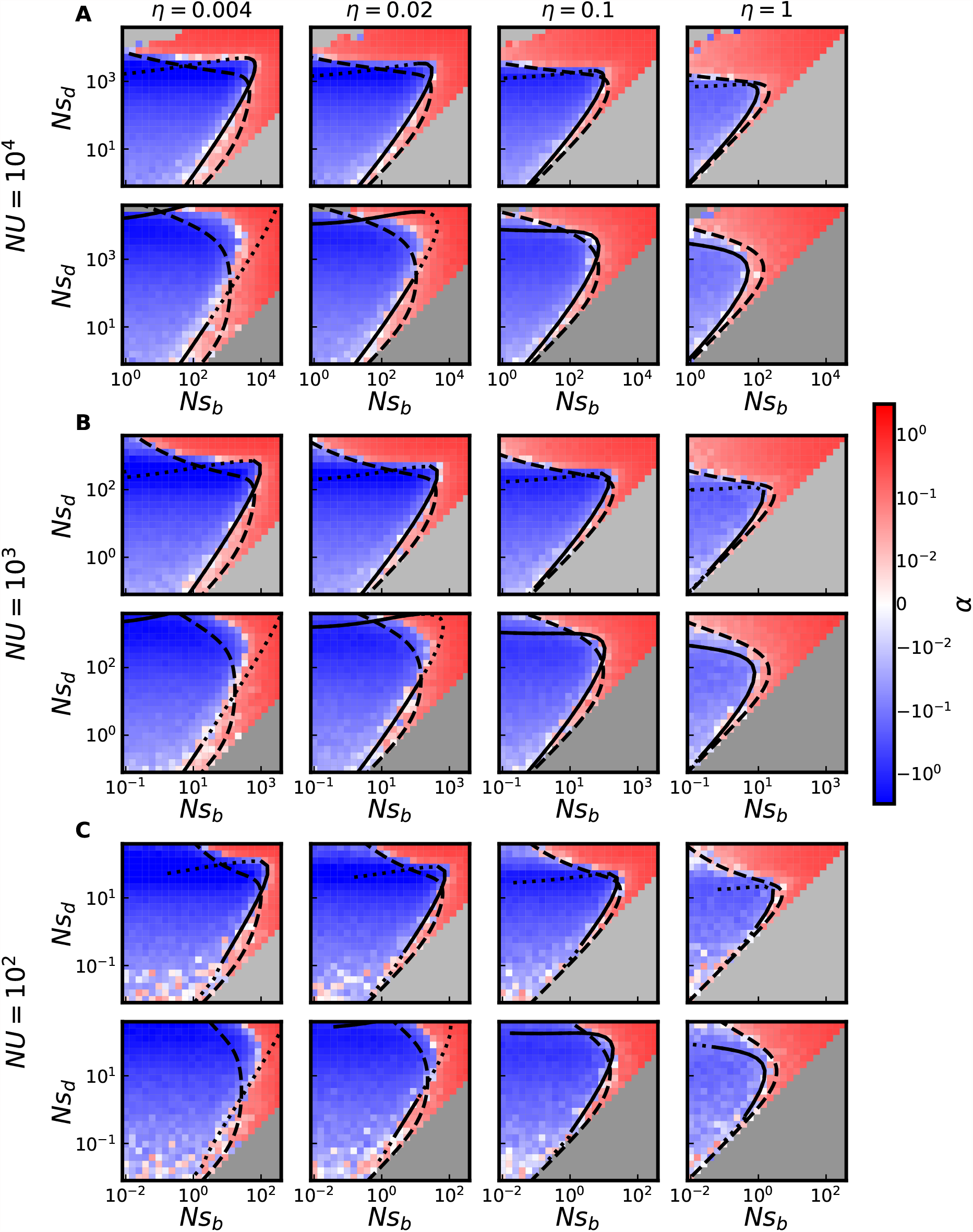
Cross sections of *α* = 0 surface as observed in simulations. Points are colored by values of *α* as observed in simulations. Values of *NU* for populations displayed in *A, B* and *C* are denoted on the left-hand side. Populations in the first, third, and fifth row are subject to two-effect DFEs; populations in the second, fourth, and sixth rows are subject to two-exponential DFEs. Solid curves denote MSSM predictions for *α* = 0 surface, with dotted lines denoting these predictions beyond the regime of validity of the MSSM approximation. Dashed lines denote predictions obtained by using our “*N*_*e*_-based heuristic” to compute *T*_*c*_ instead of the MSSM approximation.

